# Scaling down protein language modeling with MSA Pairformer

**DOI:** 10.1101/2025.08.02.668173

**Authors:** Yo Akiyama, Zhidian Zhang, Milot Mirdita, Martin Steinegger, Sergey Ovchinnikov

**Affiliations:** Department of Electrical Engineering and Computer Science, Massachusets Institute of Technology; Department of Biology, Massachusets Institute of Technology; School of Biological Sciences, Seoul National University, Seoul, Republic of Korea; Interdisciplinary Program in Bioinformatics, Seoul National University, Seoul, Republic of Korea; Institute of Molecular Biology and Genetics, Seoul National University, Seoul, Republic of Korea; Artificial Intelligence Institute, Seoul National University, Seoul, Republic of Korea

## Abstract

Recent efforts in protein language modeling have focused on scaling single-sequence models and their training data, requiring vast compute resources that limit accessibility. Although models that use multiple sequence alignments (MSA), such as MSA Transformer, offer parameter-efficient alternatives by extracting evolutionary information directly from homologous sequences rather than storing it in parameters, they generally underperform compared to single-sequence-based language due to memory inefficiencies that limit the number of sequences and averaging evolutionary signals across the MSA. We address these challenges with MSA Pairformer, a 111M parameter memory-efficient MSA-based protein language model that extracts evolutionary signals most relevant to a query sequence through bi-directional updates of sequence and pairwise representations. MSA Pairformer achieves state-of-the-art performance in unsupervised contact prediction, outperforming ESM2-15B by 6% points while using two orders of magnitude fewer parameters. In predicting contacts at protein-protein interfaces, MSA Pair-former substantially outperforms all methods with a 24% point increase over MSA Transformer. Unlike single-sequence models that deteriorate in variant effect prediction as they scale, MSA Pairformer maintains strong performance in both tasks. Ablation studies reveal triangle operations remove indirect correlations, and unlike MSA Transformer, MSA Pairformer does not hallucinate contacts after removing covariance, enabling reliable screening of interacting sequence pairs. Overall, our work presents an alternative to the current scaling paradigm in protein language modeling, enabling efficient adaptation to rapidly expanding sequence databases and opening new directions for biological discovery.

## 1 Introduction

Protein language models (pLMs) trained with self-supervision on masked amino acid prediction have demonstrated their strong applicability on a vast range of biological problems [1–4]. To improve performance, recent efforts have focused on scaling these models up to 100B parameters [5–8]. However, this scaling trend severely limits research and practical application to well-resourced industry laboratories. Interestingly, a recent study revealed that these single-sequence pLMs effectively store evolutionary statistics of protein families within their parameters, which explains the observed gains in metrics such as sequence recovery and contact prediction as models scale—larger models can store more evolutionary information at higher resolution [9]. Importantly, this very property may ultimately undermine the viability of the scaling paradigm: as sequencing data continues to grow, models will require increasingly more parameters to capture the ever-expanding sequence space, creating an unsustainable computational burden. This motivates the development of smaller but highly performant models that can adapt to new sets of sequences at inference time.

If much of the parameters in single-sequence pLMs are dedicated to storing evolutionary information, then a retrieval-based approach that uses aligned sets of homologous sequences, or multiple sequence alignments (MSAs), at inference may eliminate the need for large models. This strategy is akin to traditional statistical methods, which were developed to extract structural information from co-evolving residues [10–24]. Indeed, frontier structure prediction models such as AlphaFold2/3 rely heavily on MSAs to extract evolutionary information for accurate structure prediction [25, 26]. However, despite their groundbreaking successes in structure prediction, MSA-based approaches have not been widely adopted in self-supervised learning. MSA Transformer first demonstrated the ability of an MSA-based self-supervised deep learning model to generalize across protein families. Using only 118M parameters, it approaches the performance of ESM2-3B at long-range contact prediction [27] and outperforms it in zero-shot variant effect prediction [28].

Despite their successes, state-of-the-art MSA models, including MSA Transformer and AlphaFold3, face a fundamental limitation: they assume that all sequences in a protein family share the same structure, averaging co-evolutionary signals across the sequences of the MSA. However, subfamilies within large protein families can exhibit distinct structural and functional properties, leading to contrasting co-evolutionary patterns across phylogenetic branches. This is most evident in protein families with diverse binding interfaces or stoichiometries of self-assembled homo-oligomers [29–31]. Supporting this view, recent studies, which explore subsetting MSAs by clustering sequences or randomly masking columns of MSAs, have revealed that pockets of sequences encode co-evolutionary signals that support alternative conformations of monomeric protein structures [32–34]. These works further motivate an alternative approach to MSA-based pLMs that learn to selectively extract co-evolutionary signals most relevant to a given query sequence, rather than averaging across all sequences in the alignment.

Building on the bidirectional refinement between MSA and pair representations introduced by AlphaFold2/3, we approach these challenges by introducing MSA Pairformer, a parameter and memory efficient model that biases evolutionary signal extraction to the query sequence. Specifically, we develop the query-biased outer product operation, which selectively extracts co-evolutionary signals most relevant to the query sequence by incorporating a variant of attention aimed at maintaining high signal-to-noise with large context-windows. The model is trained via self-supervised learning to reconstruct a corrupted MSA by building both sequence and pairwise representations. MSA Pairformer outperforms models with >100x more parameters across multiple tasks, including unsupervised contact prediction, protein complex interface prediction, and zero-shot variant effect prediction. In contrast to current MSA models, MSA Pairformer extracts subfamily-specific co-evolutionary signals, identifying distinct structural properties within protein families. Lastly, through ablation studies, we investigate the role of triangle operations in disentangling direct and indirect correlations. Moreover, MSA perturbation analyses reveal that, unlike MSA Transformer, MSA Pairformer does not hallucinate contacts after removing covariance between columns of the MSA.

## 2 Related work

### Self-supervised learning of protein sequences

follows a simple procedure, wherein models are trained to recover masked amino acids from single sequences or MSAs. This approach leverages the growing set of sequencing data by enabling models to capture evolutionary statistics without explicit supervision. After training, their embeddings can be used in downstream prediction tasks, and the predicted per-residue amino acid probability distributions can be used for sequence generation, computing sequence likelihoods, and variant effect prediction [1, 7, 8, 28, 35–40]. The attention matrices of transformer-based models (e.g. ESM2 and MSA Transformer) can be used for unsupervised prediction of pairwise interactions, such as contacts in the three-dimensional structure [5, 27].

Recent efforts have focused on scaling single-sequence model sizes and their training data. These models have been shown to store evolutionary information in their parameters, making them susceptible to biases in their training data [8, 9, 41–43]. Therefore, beyond the immense computational requirements required to incorporate the rapidly growing wealth of metagenomic sequences, scaling also requires careful optimization of the training dataset. MSA models offer a promising alternative, requiring far fewer parameters to extract evolutionary information directly from alignments. Moreover, MSAs can be constructed to focus on specific protein subfamilies or filtered to mitigate phylogenetic bias, providing flexibility at inference time.

### MSA-based approaches for self-supervised learning

Despite their advantages in parameter efficiency, there are few MSA models for self-supervised learning. Most notably, MSA Transformer, which uses only 118M parameters, surpasses single-sequence models that are five times larger in unsupervised contact prediction [27]. Despite its strong performance, MSA Transformer has fundamental limitations: 1) an inability to capture subfamily-specific co-evolutionary signals; 2) high memory requirements; and 3) hallucination of contacts in the absence of covariation.

Though MSA Transformer could theoretically learn nearest-neighbor lookup via attention between sequences to compute subfamily specific conservation signal, it assumes all sequences share identical co-evolutionary patterns and ties a single attention map between positions across all sequences. This limitation is shared by AlphaFold2/3, where the outer product mean obscures subfamily-specific interactions.

The architecture also scales quadratically with sequence length and MSA depth, imposing heavy computational costs. For parameter-efficient models that extract evolutionary information at inference rather than storing it in parameters, supporting deep MSAs is crucial for providing sufficient evolutionary context. Thus, memory efficiency becomes critical to maximally benefit from rapidly expanding sequence databases. In this work, we aim to address these core limitations.

## 3 Method

### 3.1 MSA Pairformer

MSA Pairformer addresses the limitations of both large-scale single-sequence models and existing MSA models through a parameter-efficient architecture that learns to extract evolutionary signal most relevant to a query sequence. Using only 111M parameters, MSA Pairformer processes MSAs through alternating updates to MSA and pair representations across 22 layers (Figure 1A). The model takes as input an MSA, where rows correspond to homologous sequences and columns correspond to residue positions. Each layer performs three sequential operations: (1) pair-weighted averaging updates the MSA representation using pairwise dependencies stored in the pair representation, (2) query-biased outer product updates the pair representation using the MSA representation by focusing on sequences most relevant to the query, and (3) triangle multiplicative updates refine the pair representation by considering triplet interactions between residues. This architecture builds off of AlphaFold3’s MSA module, which is specifically trained for supervised structure prediction. While self-supervised models to date do not explicitly build pairwise representations and rely on extracting pairwise relationships post-hoc from attention weights, we hypothesize that modeling pairwise relationships throughout training may encourage the model to decompose complex co-evolutionary signals, as well as improve the sequence representations.

**Figure 1:**
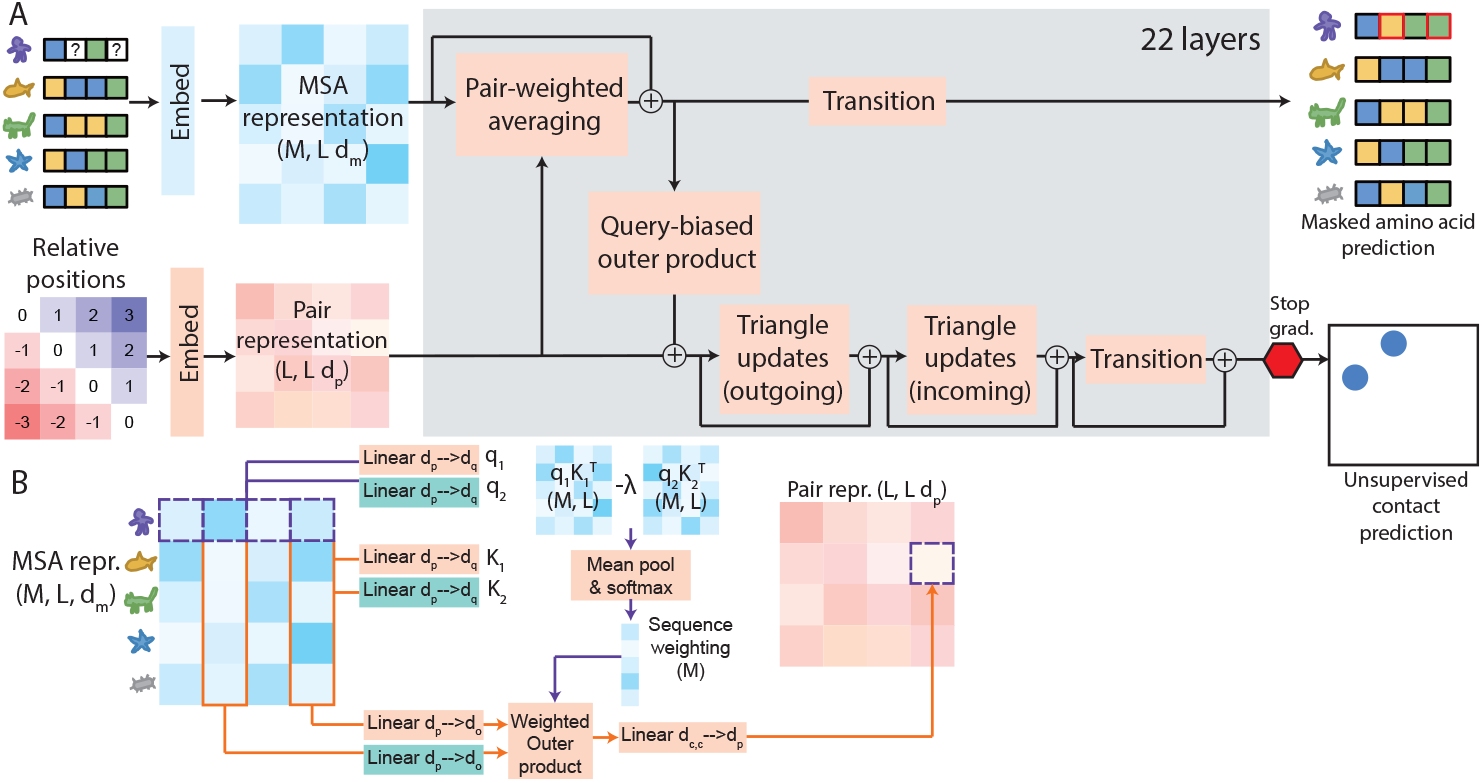
MSA Pairformer architecture. A) Overview showing the integration of MSA representation (blue) and pair representation (orange) across 22 layers. Each layer performs three operations: pair-weighted averaging, query-biased outer product, and triangle updates. The model is trained on masked amino acid prediction and enables unsupervised contact prediction from pair representations. B) Detailed view of the query-biased outer product block, showing pre-softmax differential attention computation that weighs sequences based on their evolutionary relevance to the query.

#### 3.1.1 Query-biased outer product

A key innovation of MSA Pairformer is its query-biased approach to extracting evolutionary information (Figure 1B). Rather than averaging co-evolutionary signals across all sequences equally (as in prior MSA models), MSA Pairformer learns to weight sequences based on their evolutionary relevance to the query sequence. This addresses the limitation that protein subfamilies within large families often exhibit distinct co-evolutionary patterns.

For each sequence *s*, the model computes an attention weight *a*_*s*_ that measures the evolutionary relevance of *s* to the query sequence. These attention weights determine the contribution of each sequence to the pair representation update:

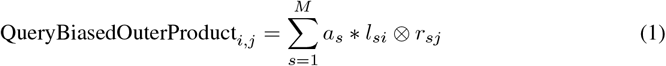

where *l*_*si*_ and *r*_*sj*_ are learned projections of the MSA representations for sequence *s* at positions *I* and *j*. In order to compute *a*_*s*_, MSA Pairformer uses pre-softmax differential attention.

**Pre-softmax differential attention** builds off of several studies, primarily, the Differential Transformer, and aims to reduce noise in attention weights and prevent attention dispersion that occurs with long contexts in softmax attention [44–46]. Unlike differential attention, which computes differences after softmax normalization, pre-softmax differential attention computes the differential prior to softmax:

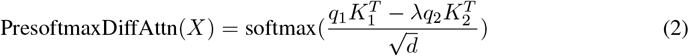

where *λ* is a learned scaling factor and reparameterized as in Differential Transformer [45], and *q*_1_, *q*_2_, *K*_1_, and *K*_2_ are learned query and key projections of the sequence representations.

To obtain sequence-level attention weights *a*_*s*_, the model first computes position-wise pre-softmax differential attention scores between the query sequence and each sequence *s* at every position *i*. These position-wise scores are then averaged across all positions and normalized with softmax.

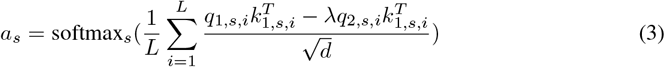

We hypothesize that this approach may enable the model to tune attention sharpness while preserving the stability and generalization benefits of QK-Norm and weight decay. We compare this approach against standard attention, differential attention, and a non-negative variant of differential attention. We find that the pre-softmax differential improves masked amino acid recovery in MSA Pairformer (full details in Appendix).

### 3.2 Pretraining, attention finetuning, and unsupervised contact prediction

We divide training into two phases: pre-training and attention finetuning. For both, we use the 270K Uniclust30 MSAs from the OpenProteinSet, using the same train and validation splits for each phase [47]. In the pre-training phase, we use uniform sequence weighting in the outer product layers. In the attention finetuning phase, we replace the outer product mean with query-biased outer products.

Lastly, we train a logistic regression classifier, which takes as input the 256-dimensional pair representation and predicts residue contacts. We define contacts as residue pairs with C*β*-C*β* < 8Å. For glycines, the C*β* is placed based on the N, CA, C vector of the backbone, following the protocol of TrRosetta [48]. We use the same 20 training samples used to train the MSA Transformer and ESM-family contact heads and fit a logistic regression contact head for each layer in MSA Pairformer. The MSAs and contacts for these 20 samples are derived from the trRosetta training set (full details provided in Appendix).

## 4 Results

### 4.1 State-of-the-art unsupervised contact prediction using two orders of magnitude fewer parameters

Unsupervised contact prediction is widely used to evaluate the ability of a self-supervised pLM to capture long-range dependencies between residues. Here, we compare MSA Pairformer’s performance on long-range contact precision (*≥* 24 residue separation) to that of MSA Transformer and the ESM2-family models for CASP15 targets. For fair comparison with the ESM2 models, which were trained on UniRef sequences, we generated MSAs using hhblits to search the UniRef30 (ver. Feb 2023) [49]. We standardized the experimental conditions by limiting MSA depth to 512 sequences (the maximum depth MSA Transformer can process for these targets due to memory constraints) and restricting evaluation to the 45 proteins shorter than 1024 residues (MSA Transformer’s maximum length). Full dataset metrics are provided excluding only T1164, which at 3364 residues exceeds the memory capacity of these models. We do not compare to ESM C, since they lack a publicly available contact prediction head and are trained on metagenomic sequences.

Among ESM2-family single-sequence models, long-range contact precision scales with model size, peaking at an average P@L of 0.46 for ESM-15B (Fig 2A). MSA Transformer achieved an average P@L of 0.44, outperforming ESM-650M but falling short of ESM-3B. Remarkably, MSA Pairformer substantially outperformed all baselines, achieving an average P@L of 0.52, a 6% point improvement over ESM-15B despite a more than 100x reduction in parameter count (Fig 2A, B). This represents an 8% point improvement over MSA Transformer (0.52 vs. 0.44), highlighting the effectiveness of the Pairformer architecture for modeling residue-residue interactions.

**Figure 2:**
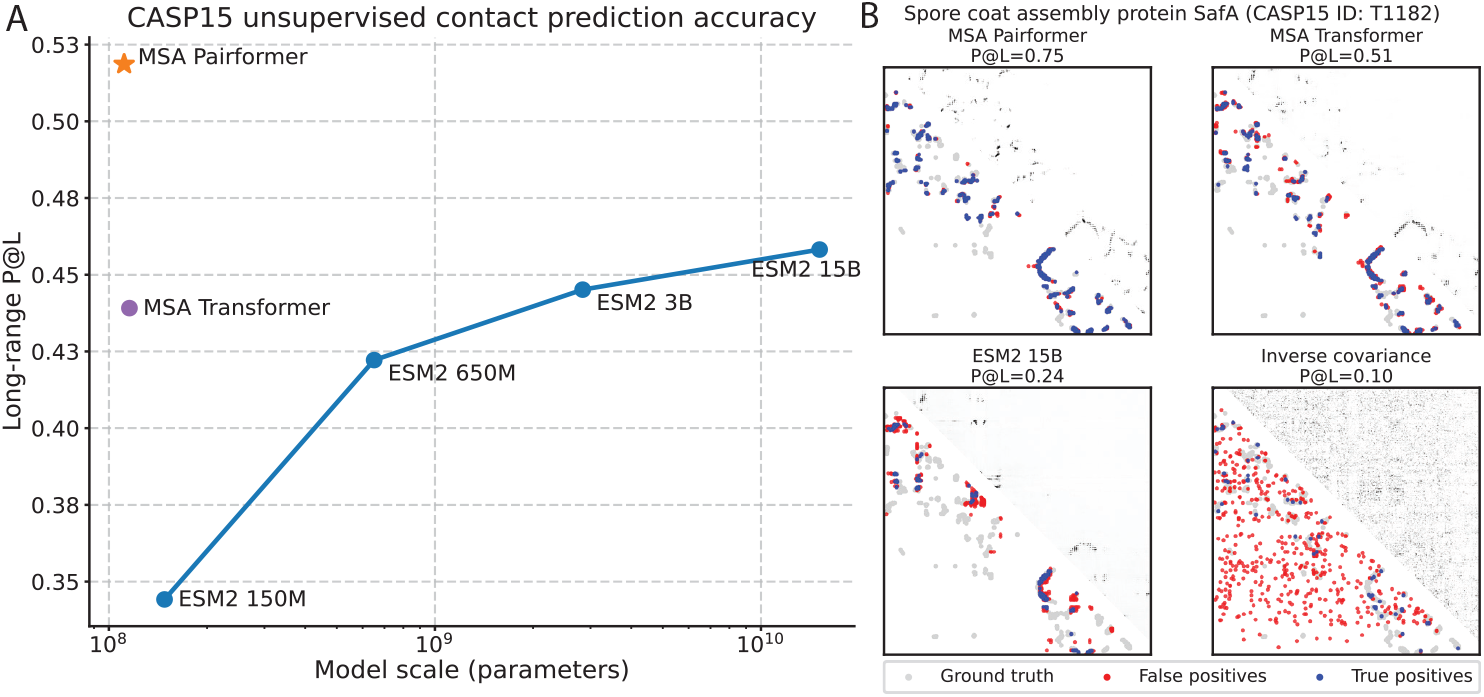
MSA Pairformer achieves state-of-the-art contact precision on CASP15 using two orders of magnitude fewer parameters. (A) Long-range contact precision at L (P@L, where L is sequence length) across model sizes (B) Long-range contact prediction for T1182. Grey points indicate all ground truth long-range (*≥* 24 residue separation) contacts from crystal structures; red and blue points show true and false positives predicted by each method.

Interestingly, unlike MSA Transformer and ESM2 models, MSA Pairformer does not benefit from average product correction (APC). APC was introduced in statistical methods for contact prediction to remove background phylogenetic signal that creates spurious correlations unrelated to physical contacts. These results suggest that MSA Pairformer’s pair representations distinctly decompose phylogenetic signals from contacts.

We also find that using query-biased attention increases MSA Pairformer’s contact precision from 0.50 to 0.52. This supports the notion that extracting the co-evolutionary signal most relevant to the query sequence is beneficial to building the most informative representations.

### 4.2 Accurate prediction of protein-protein interactions

Beyond monomeric structures, accurate prediction of interactions between different proteins (i.e. hetero-oligomeric interactions) is crucial for understanding how proteins function in complexes. We therefore evaluated MSA Pairformer’s ability to predict contacts at the interface of protein-protein interactions using paired MSAs of 25 evolutionarily conserved protein complexes from Ovchinnikov et al. [13], comprising 31 interacting proteins. We computed the precision of the top *K* predicted contacts (P@K), where *K* equals the total number of interface contacts in the crystallographic structure. We compare to MSA Transformer, ESM2-15B, and gLM2 [50]. For direct comparison with MSA Transformer, we excluded two complexes (three interactions) exceeding the 1024 residue limit.

MSA Pairformer significantly outperforms all baseline methods (Mann-Whitney U test adj. p-value *≤* 0.05), achieving a median P@K of 0.53 compared to MSA Transformer’s 0.29, ESM2-15B’s 0.01, and gLM2’s 0.01 (Figure 3A). MSA Pairformer consistently outperforms MSA Transformer on all but two targets, where performance is roughly equivalent (Figure 3B). This nearly two-fold improvement over MSA Transformer demonstrates MSA Pairformer’s substantial improvements in identifying protein-protein interface contacts. In general, we find that MSA Transformer greatly suppresses interface contacts. For MSA Pairformer, the median predicted probabilities of correctly predicted contacts decreases 1.4-fold from 0.85 to 0.61 for monomeric and interface contacts, respectively. By comparison, MSA Transformer shows a 4.4-fold reduction, dropping from 0.69 to 0.15 (Figure 3C). Moreover, we find that high-confidence predictions of interacting residues from MSA Pairformer are almost always accurate. Among 478 predicted contacts with probabilities greater than 0.8, 85% are within 8 and 95% are within 12 in the crystallographic structure (Figure 3D). Interestingly, in some cases, MSA Transformer entirely suppresses interface contacts despite strong co-evolutionary signal, as evidenced by inverse covariance analysis (Figure 3E).

**Figure 3:**
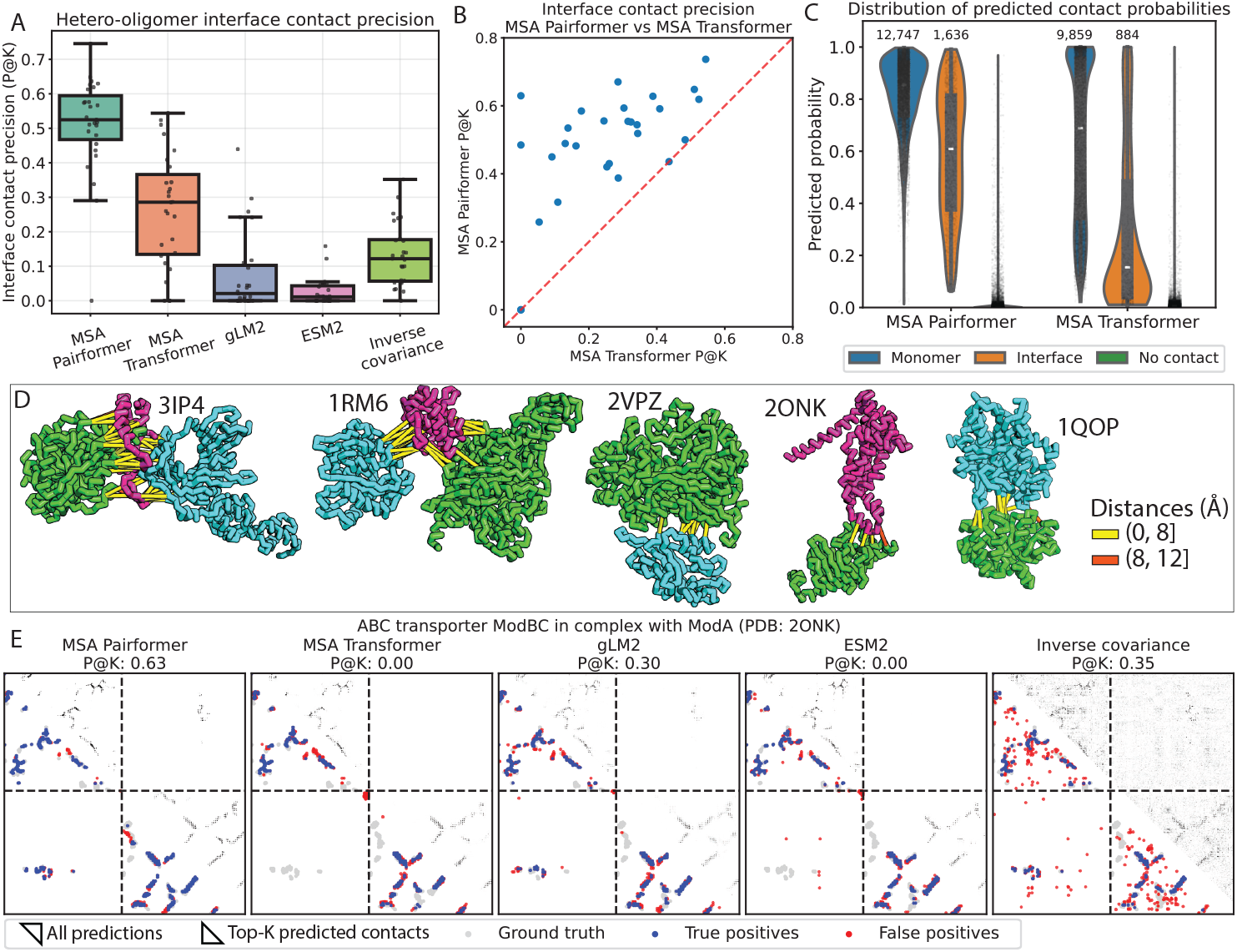
MSA Pairformer substantially outperforms all methods in predicting residue-residue interactions between hetero-oligomeric complexes. A) Box plot comparing the complex interface contact precision, P@K, where K is the total number of interface contacts in the ground truth structure. B) Head-to-head comparison of MSA Pairformer and MSA Transformer predictions. C) Distribution of predicted contact probabilities for [blue] correctly predicted top-L monomer contacts, [orange] correctly predicted top-K interface contacts and [green] 100,000 randomly selected pairs of non-interacting residue pairs. Number above each distribution indicates the number of correctly predicted contacts in each class. D) Crystallographic structures for six examples in the dataset. Yellow and orange bars indicate predicted contact probabilities *≥* 0.8. Predicted contacts that are within 8 or 12 in the crystal structure are colored in yellow and orange, respectively. The structures are pulled apart for clarity. C) Contact predictions for the ABC transporter ModBC and ModA (PDB: 2ONK). Lower triangle shows (grey) ground truth, (blue) true positive and (red) false positive contacts, using the top-L for each monomer and top-K for the interface, where *L* is the length of the monomer and *K* is the number of interface contacts in the crystallographic structure. Upper triangle shows all predictions as a heatmap; darker color indicates higher predicted contact probability.

Our evaluations reveal that single-sequence models particularly struggle to predict interface contacts. While ESM2 was only trained using individual protein sequences in isolation, interacting proteins are often encoded in a single fusion gene across the tree of life. Indeed, 15 of the 31 interactions in this evaluation set involve proteins encoded as fusions in subsets of organisms, meaning that ESM2 trained on sequences containing both proteins. However, ESM2 still fails to accurately predict contacts for these examples. Furthermore, despite training on metagenomic contigs spanning interacting proteins, gLM2 was generally outperformed by the inverse covariance baseline. These results reveal potential fundamental limitations of current single-sequence models that extend beyond training data composition.

### 4.3 Extracting subfamily-specific structural properties

We next evaluated whether query-biased outer products enable MSA Pairformer to extract structural properties that drastically differ between subfamilies. We focused on the bacterial response regulator protein family, a widely used case study for demonstrating a method’s ability to disentangle subfamily-specific co-evolutionary signals [9, 31, 51]. Experimental structures (Figure 4A) are available for representatives of three distinct subfamilies, OmpR, LytTR, and GerE, all of which form homodimers composed of two identical protein chains [52]. These structures share similar intra-chain contacts within each monomer (Figure 4A) but exhibit divergent inter-chain contacts at their homodimeric interfaces (Figure 4A).

**Figure 4:**
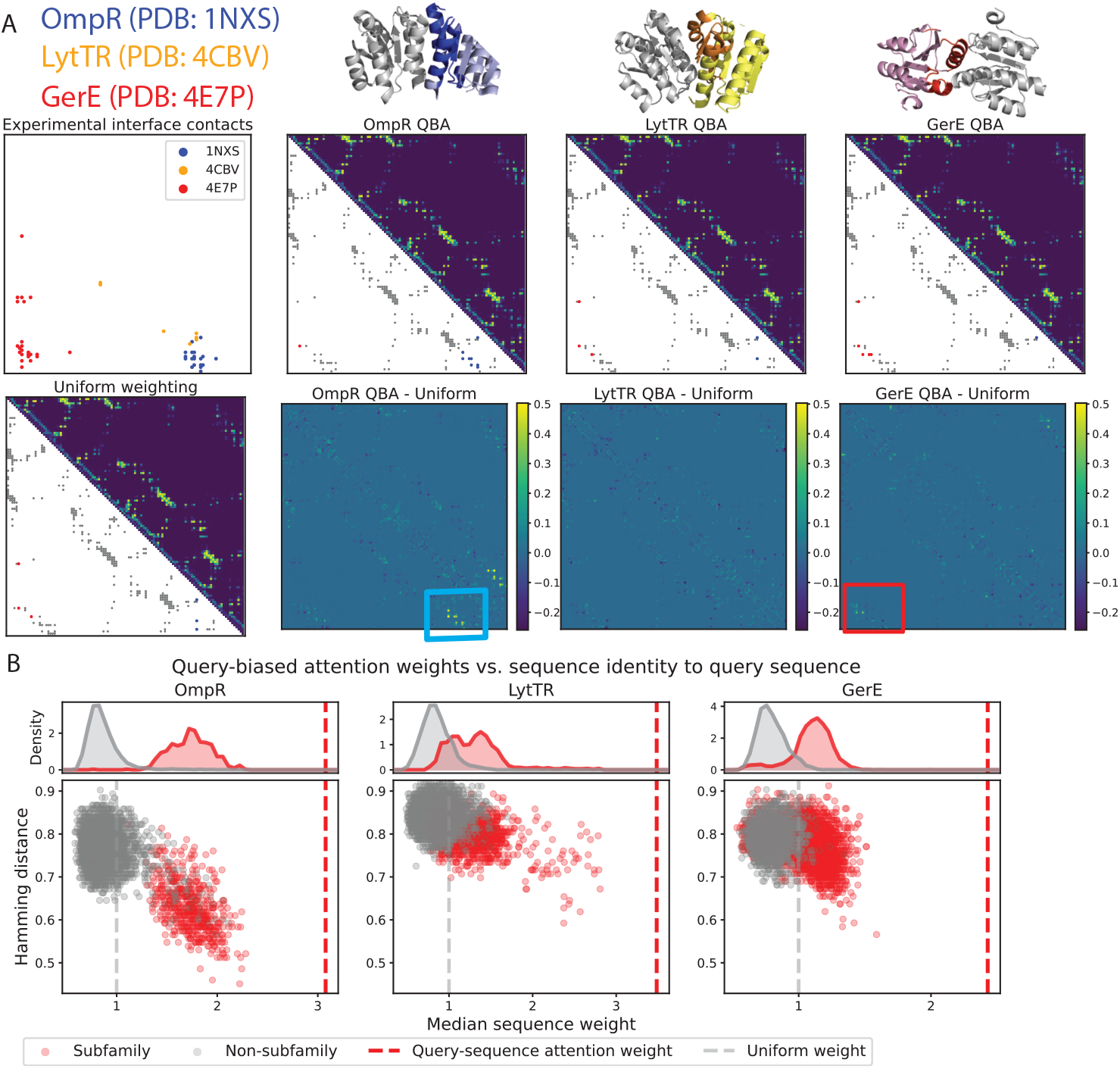
Query-biased outer product enables MSA Transformer to extract subfamily-specific homo-oligomeric contacts. A) Experimental dimeric structures for representatives of three response regulator (RR) subfamilies: OmpR (PDB ID: 1NXS), LytTR (PDB ID: 4CBV), and GerE (PDB ID: 4E7P). One chain is highlighted in color to show the homodimer interface. [Top row] shows subfamily-specific inter-chain contacts (residue pairs *≥* 6 residue separation) and predictions using MSA Pairformer with query-biased outer product. Upper triangle shows the predicted contact probabilities, and the lower triangle shows the top contact predictions, with colored dots indicating known subfamily-specific inter-chain contacts. [Bottom row] shows difference in predicted probabilities when using query-biased outer product versus uniform sequence weighting. B) Median sequence weight across the layers of the model versus Hamming distance to the query sequence. Top panels show distribution of sequence attention weights for subfamily members (red) and non-subfamily sequences (grey). Grey dotted line indicates weights used for uniform sequence attention and red dotted line indicates weight assigned to the query sequence.

Using a single MSA with 4096 sequences comprised of 2506, 1074, and 516 sequences from the GerE, LytTR, and OmpR subfamilies, respectively, we predicted contacts for the three subfamilies by setting the query sequence as the one used in the representative crystal structures. For each prediction, we selected the top *N* contacts, where *N* is the total number of contacts in the experimental structure, and evaluated the recovery of subfamily-specific homodimeric interface contacts (Figure 4A). Query-biased outer product had the strongest effect on recovering OmpR subfamily-specific homodimeric contacts, recovering 8/19 subfamily-specific interface contacts compared to only 3/19 with uniform sequence weighting. This can be explained by the fact that OmpR sequences make up only 12% of this MSA, and therefore, without query-biased attention the co-evolutionary signals distinct to this subfamily are averaged out. Nonetheless, even for the GerE subfamily, which makes up the dominant 61% of sequences in the MSA, query-biased outer product improves predictions, identifying 5/23 interface contacts versus 3/23 with uniform weighting. The single OmpR interface contact predicted using uniform sequence weighting decreased in contact probability by 14% points from 0.63 to 0.46 when the GerE sequence is assigned as the query. These results suggest that MSA Pairformer’s query-biased outer product is crucial for extracting subfamily-specific co-evolutionary signals.

Interestingly, we observe bimodal distributions in the median sequence weight assignments for each sequence across subfamily predictions, where subfamily members are significantly upweighted compared to non-subfamily sequences (Mann-Whitney U p-value < 0.05; Figure4B). For OmpR, which represents a simpler case since most OmpR sequences in this MSA have higher sequence identity to the OmpR query, all of the 516 subfamily sequences are upweighted relative to uniform weighting. Strikingly, for the GerE subfamily sequences, 81% of the subfamily sequences were upweighted compared to uniform weighting, including sequences with as low as 9% sequence identity. This result highlights MSA Pairformer’s ability to assign biologically relevant sequence weights across an MSA.

Overall, these results demonstrate that query-biased attention enables MSA Pairformer to better capture subfamily-specific structural properties. This capability is biologically significant, as subfamily-specific structural differences often underlie distinct signaling mechanisms, regulatory functions, and protein-protein interactions that are central to understanding protein function and evolution.

### 4.4 MSA Pairformer overcomes the contact-variant effect prediction trade-off in protein language models

While scaling single-sequence models results in better unsupervised contact prediction, the opposite has been found in variant effect prediction [28, 8]. Indeed, in ProteinGym zero-shot variant effect prediction evaluations, the performance of ESM2 models peaks with 650M parameters, decreases for both 3B and 15B. Here, we sought to evaluate whether MSA Pairformer’s state-of-the-art contact prediction performance comes at the cost of poor zero-shot variant effect prediction by benchmarking on the 219 ProteinGym deep mutational scan substitution experiments [28].

Although studies have shown the benefits of carefully constructing MSAs, ensembling predicted amino acid distributions using many MSAs, and ensembling predicted distributions from multiple versions of the model, we simply use the MSAs provided by the ProteinGym benchmarking team [28, 53–55]. Strong performance on a single, unrefined MSA would highlight MSA Pairformer’s robustness to potentially adversarial sequences in the MSA. For each deep mutational scan experiment, we sample up to 4096 sequences from the alignment, weighting the probability of selecting a sequence by the inverse of the number of sequences it is at least 80% identical to and use a maximum sequence length context window of 896 residues.

MSA Pairformer substantially outperforms all ESM2 and ESM C models, as well as the MSA Transformer ensemble predictions, achieving an average Spearman’s correlation of 0.47 (Figure 5A). This represents a 4% increase compared to MSA Transformer, and an 11% increase over ESM2 15B. Importantly, whereas the performance in variant effect prediction decreases as single-sequence pLMs improve in contact prediction, MSA Pairformer uniquely achieves strong performance in both contact prediction and variant effect prediction (Figure 5B).

**Figure 5:**
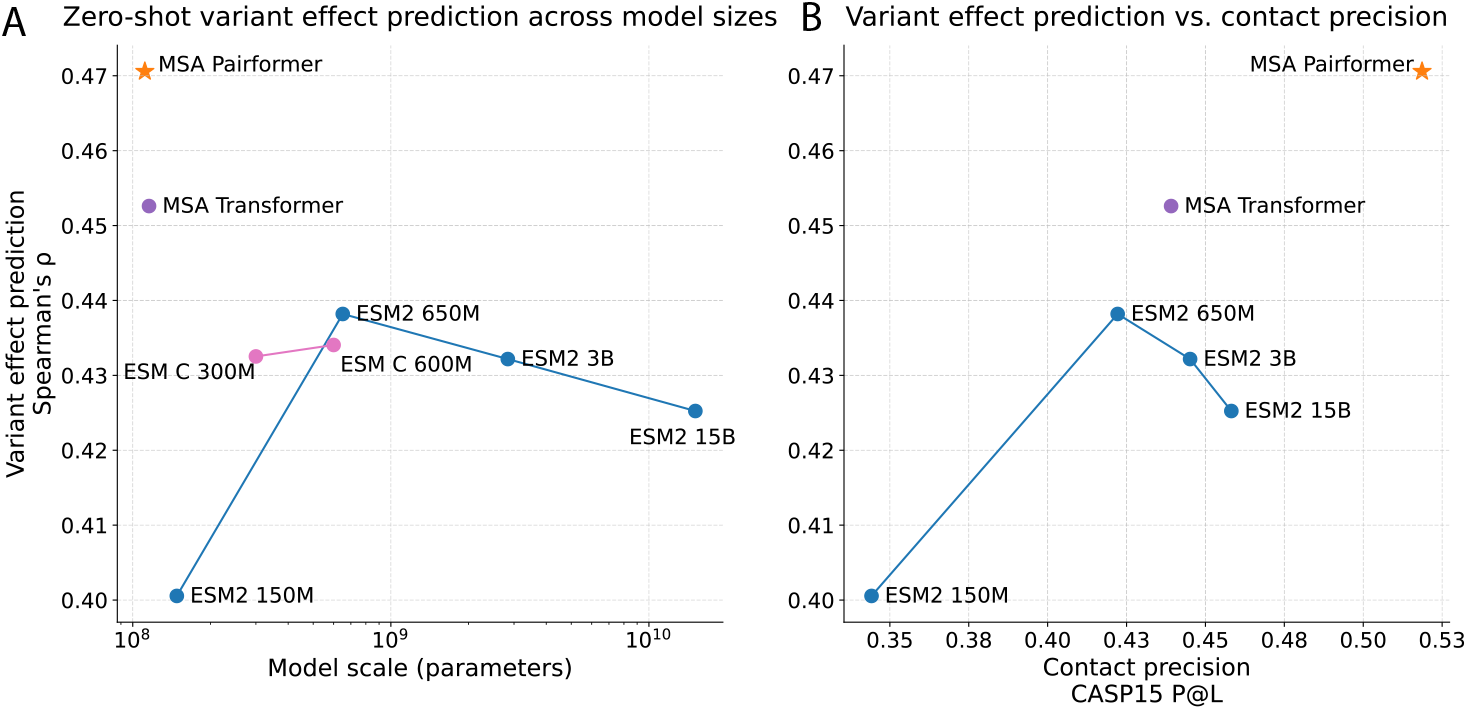
MSA Pairformer excels at both contact and zero-shot variant effect prediction, pushing beyond the scaling wall exhibited by single-sequence models. ProteinGym zero-shot variant effect prediction results A) across model sizes, and B) compared to contact precision (CASP15 long-range P@L). MSA Pairformer achieves state-of-the-art contact accuracy performance, while maintaining strong variant effect prediction.

### 4.5 MSA Pairformer pseudolikelihoods better differentiate binding and non-binding PPI interface sequeunces

Rationally reprogramming protein-protein interactions in two-component pathways is a longstanding challenge in the design of synthetic signaling circuits. Applying pLMs to coordinate sequence design of PPI interfaces remains unreliable due to their inability to model hetero-oligomeric interactions. Given that MSA Pairformer accurately models PPIs, we hypothesized that its pseudolikelihood scores could be used to discriminate binding and non-binding pairs of sequences. Using a library of mutants at four key interface residues in the ParD3 antitoxin from Aakre et al. [56], we evaluate the correspondence between model-assigned pseudolikelihood and toxin-antitoxin binding. This dataset contains a 9194 sequences, of which 252 are good binders (average fitness *≥* 0.5). We compute pseudolikelihoods using the four mutated positions to rank the sequences and compute the precision of the top 252 sequences. MSA Pairformer achieves a precision of 0.58, outperforming MSA Transformer (0.50) and ESM2-15B (0.39) by 8% points and 19% points, respectively (Figure 6A, B). These results are consistent with their ability to capture residue-residue interactions at the toxin-antitoxin interface (Figure 6C, D). In order to test whether MSA Pairformer’s performance gains derive from its ability to model the ParD3-ParE3 interaction, we recompute scores using only ParD3 sequences. We find that its performance deteriorates sharply to 0.45, approaching that of ESM2. These results suggest that modeling the ParD3-ParE3 interaction is integral to the improved agreement between MSA Pairformer’s pseudolikelihood scores and toxin-antitoxin binding.

**Figure 6:**
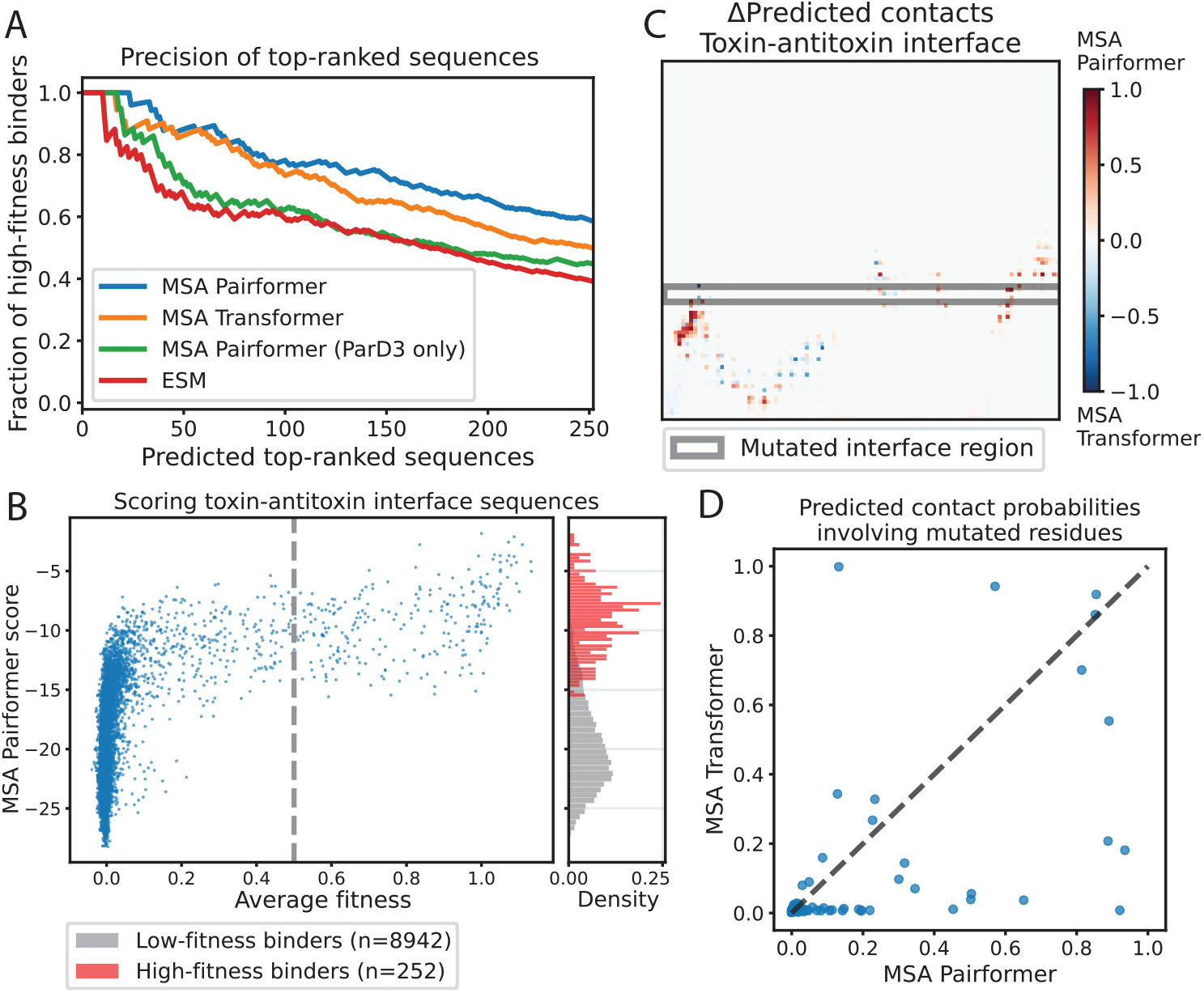
MSA Pairformer pseudolikelihood scores better discriminate binding and non-binding PPI interface sequences. A) Precision of the top-ranked sequences. Line positions indicate the fraction of high-fitness binders (y-axis) among the top-K ranked sequences assigned by model pseudolikelihood scores (x-axis). B) MSA Pairformer pseudolikelihood scores versus average fitness for every sequence. Dotted line shows 0.5 average fitness score boundary. Histograms in the rightmost pannel show normalized distributions of MSA Pairformer scores for high-fitness binders (red) and low-fitness binders (grey). C) Difference in predicted contact probabilities between MSA Pairformer and MSA Transformer for the ParD3-ParE3 toxin-antitoxin interface. Grey box indicates the areas overlapping the four mutated ParD3 residues. D) Scatterplot of contact probabilities for interface residue pairs involving the four mutated ParD3 residues.

### 4.6 Exploring potential functions of triangle multiplicative updates via ablation studies

Given MSA Pairformer’s particularly strong performance in contact prediction, we sought to better understand its key underlying mechanisms. Triangle Multiplicative Updates were first introduced in AlphaFold2, alongside Triangle Attention, as part of the triangular operations framework. These operations are designed to incorporate the triangle inequality as an inductive bias, encouraging the model to consider the geometric constraints that govern spatial relationships between residue triplets. Although previous studies have demonstrated the importance of these triangle updates for accurate structure prediction [25, 57], the specific mechanisms underlying their effectiveness remain poorly understood. To gain insights into their potential function, we performed an ablation study on the triangle multiplicative updates.

In proposing a potential mechanism for triangle updates in MSA Pairformer, we draw inspiration from traditional methods for extracting co-evolutionary statistics from MSAs, such as direct coupling analysis, inverse covariance, Potts Hamiltonian models and GREMLIN [10–24]. These approaches rely on disentangling direct and indirect correlations between residue positions. For example, the off-diagonal elements of the inverse covariance matrix (or precision matrix) are negatively proportional to the partial correlation between two positions of a protein. That is, given an inverse covariance matrix Ω = Σ^*−*1^, the partial correlation *ρ*_*i,j*_ between positions *i* and *j* can be calculated as 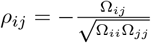. This is also equivalent to computing the correlation between the residuals *e*_*i*_ and *e*_*j*_ after regressing out effects from all other residues.

Given that triangle updates consider the pair representations *z*_*ik*_ and *z*_*jk*_ for all residues *k* when processing the pair representation *z*_*ij*_, we hypothesize that one of its potential roles in MSA Pairformer is to similarly remove the effects of indirect correlations from the pair representation (Figure 7A). To explore this hypothesis, we remove the model’s ability to consider residue triplets. Rather than summing the Hadamard product of the representations *a*_*i,k*_ and *b*_*j,k*_ across all residues *k*, we limit the update to interactions between (*i, j*) and (*j, i*) (Algorithm 1, Figure 7A). This ablation maintains the total parameter count while preventing the model from considering potential shared interactions with other residues when updating a given pair. We refer to this ablation as Pair Updates. By removing triplet processing entirely, Pair Updates represents a more drastic ablation than previous studies that replaced triangle operations with biaxial attention, which preserves some triplet information through two of the three triangle edges. Pair Updates may be more similar to traditional mutual information approaches, where *a*_*ii*_ and *b*_*jj*_ represent marginal distributions, while *a*_*ji*_ and *b*_*ji*_ are analogous to joint distributions.

**Figure 7:**
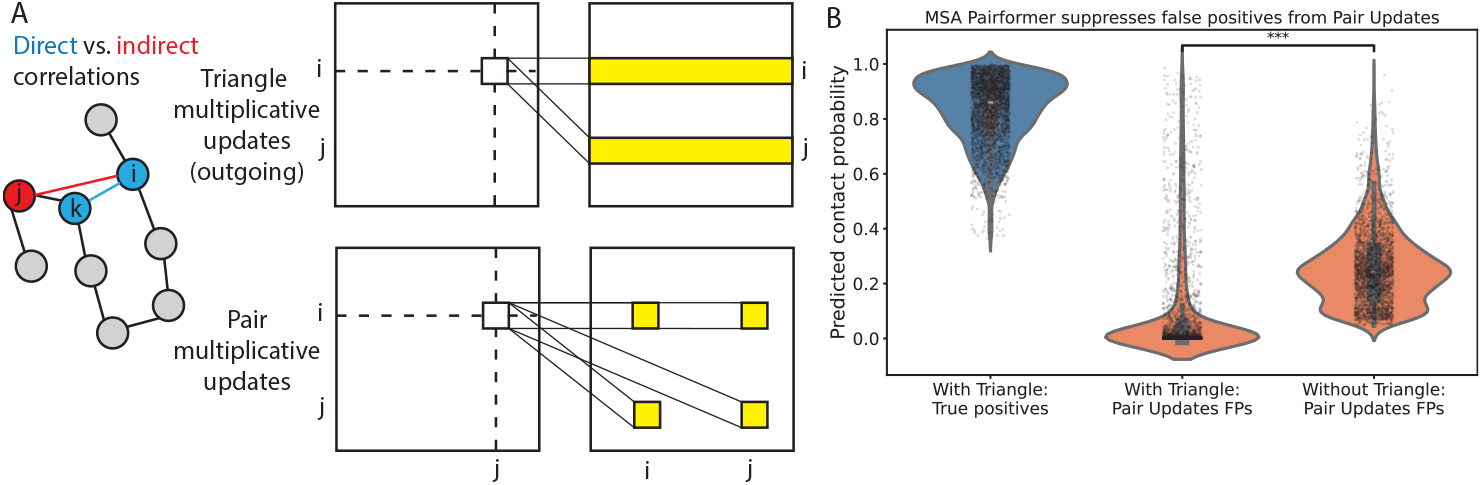
Triangular operations enable MSA Pairformer to disentangle direct and indirect correlations. A) Illustration comparing direct and indirect correlations in protein structures. Grids illustrate triangle updates (top) and pair updates (bottom) applied to MSA Pairformer’s pairwise representation. B) Distributions of probabilities assigned by MSA Pairformer with triangle updates (left and middle) and without triangle updates (right). [Left]: probabilities of true positives from MSA Pairformer; [center] probabilities assigned by MSA Pairformer to false positives from Pair Updates; and [right] probabilities assigned by Pair Updates to false positives from Pair Updates

#### Algorithm 1

Triangular / Pair multiplicative update using “outgoing” edges

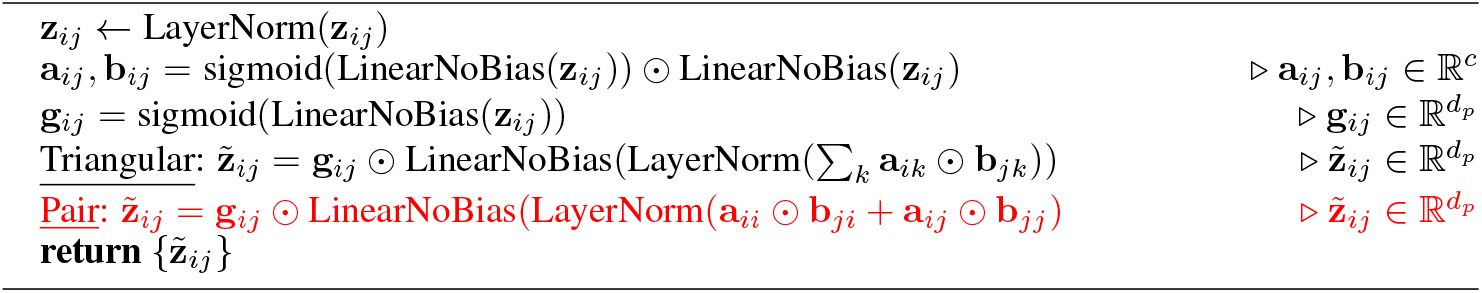

We initialize the Pair Updates model using the base pretrained model and fine-tune for 12,500 steps. We then fit contact heads to layers 15-21. As before, layer 15 showed the highest validation P@L. Interestingly, despite only a minor increase in test set perplexity (from 3.62 to 3.97), we find that long-range P@L on the CASP15 targets decreased from 52% to just 34%.

In order to test whether this decrease in contact accuracy is due to the Pair Updates model’s inability to remove indirect correlations, we focus on 16 CASP15 targets for which MSA Pairformer with triangle updates achieves at least 75% long-range P@L. Using these examples, we compute the fraction of the false positives from Pair Updates could be explained by a mediating residue. That is, for each false positive contact prediction (*i, j*), we checked whether there exists a residue *k* such that both (*i, k*) and (*j, k*) are true contacts, which would suggest that the model incorrectly inferred a direct (*i, j*) contact from their shared interaction with *k*. We refer to the fraction of these explainable false positives as the mediation rate. We then test whether false positives are enriched for indirect contacts compared to randomly sampled pairs. For each (*i, j*) false positive pair, we randomly select a residue *p* that is at least 24 residues from *i* and repeat the triplet analysis. If no residue *k* is found such that (*i, k*) and (*p, k*) are true contacts, we repeat the analysis by randomly selecting some residue *p* that is at least 24 residues from *j*. We run this analysis 1,000 times to build a null distribution of mediation rates. Interestingly, we find that 12 out of 16 targets were significantly enriched in false positive pairs with high mediation rates (adj. p-value *≤* 0.05). The four targets with no significant enrichment were among the five shortest proteins in the set, which is likely explained by the greater overlap in random long-range pairs and false negatives for shorter proteins. On average, we find that 41% of false positives from Pair Updates can be explained by a third residue for which the pair of residues are both in contact. As a control, we repeat this analysis using MSA Pairformer predictions that were randomly noised until the long-range P@L approximated the precision of Pair Updates. Randomly noising the MSA Pairformer predictions allows us to establish a separate baseline which should represent randomly poor predictions. Indeed, the false positives from these noised MSA Pairformer predictions are not significantly enriched in high mediation rates compared to their null distributions (all adj. p-value ≤ 0.05), further suggesting that false positives after ablating triangle updates are enriched in indirect correlations. Lastly, we find that MSA Pairformer (using triangle updates) assigns significantly lower probabilities to these false positive pairs (Mann Whitney U p-value *<* 0.05; Figure 7B). These results support the notion that ablating triangle updates impairs MSA Pairformer’s ability to account for indirect correlations.

The analysis above suggests potential roles of triangle updates in the model that was trained with triangle updates, but is insufficient to suggest whether triangle updates are necessary for the model’s ability to properly account for indirect correlations. We therefore trained a pair updates model from scratch and trained a contact prediction head, comparing to a baseline triangle updates model using the same hyperparameters. For this experiment, we train and evaluate both models without query-biased outer product for simplicity. On a held out set of 400 MSAs, the triangle updates model slightly outperforms the pair updates model in masked amino acid prediction, achieving average perplexities of 3.98 versus 4.12 and average accuracies of 0.59 versus 0.58 for the triangle and pair models, respectively. However, the pair updates model performs substantially worse in contact prediction, resulting in a 16% point decrease in long-range P@L (0.36 vs. 0.52), supporting the notion that training MSA Pairformer with triangle updates is critical for its strong performance in unsupervised contact prediction.

Although these analyses do not provide a complete algorithmic understanding, the results support our hypothesis: by considering the interactions of all possible triplets of residues when building pair representations, triangle updates may play a crucial role in disentangling direct and indirect correlations encoded in the MSA’s covariance statistics.

### 4.7 MSA perturbations reveal distinct mechanisms between MSA Pairformer and MSA Transformer

To further investigate how MSA Pairformer and MSA Transformer extract pairwise dependencies, we analyzed their responses to MSA perturbations. Rao et al. [27] demonstrated a peculiar behavior with MSA Transformer: the model often maintains high contact prediction accuracy even when covariance signals are ablated, suggesting it may rely on mechanisms beyond co-evolutionary signal extraction.

We tested how the two models respond to the two types of MSA perturbations introduced by Rao et al. [27]:

1. Shuffled covariance: randomly permuting values within each column of the MSA. This preserves individual position profiles (amino acid frequencies) but removes pairwise correlations between columns.
2. Shuffled positions: permuting the order of columns. This maintains pairwise correlations but tests whether the model is sensitive to the sequential arrangement of positions.

Both perturbations should substantially reduce contact prediction accuracy in a model that properly extracts evolutionary information. Evolution constrains amino acid identities jointly across positions, and sequences from permuted MSAs would generally not fold into the original structure. Therefore, an MSA model that robustly captures evolutionary signals should not predict contacts of the original structure after covariance is ablated. Furthermore, permuting the order of amino acids in a protein sequence will generally not create a well-structured protein, and thus a model that considers physical constraints should not predict contacts from the original structure for such sequences.

Using 96 randomly selected targets from the trRosetta training set (maximum 512 residues), we evaluated both MSA Pairformer and MSA Transformer under these perturbations (Table 1, Figure 8). Both models show substantial deterioration in contact accuracy when positions are shuffled, indicating that they consider the sequential organization of amino acids, rather than simply extracting pairwise correlations.

**Table 1:**
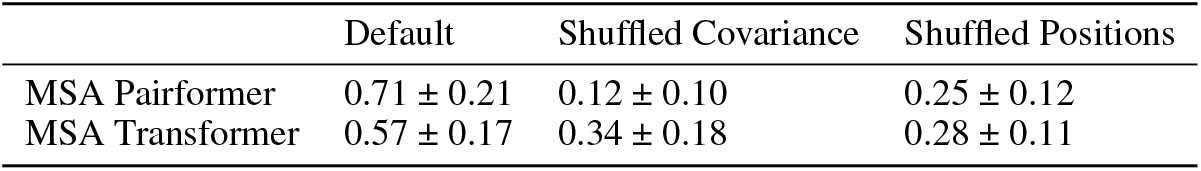
MSA Pairformer appropriately fails to predict contacts when co-evolutionary signal is destroyed, while MSA Transformer hallucinates similar contact patterns despite the absence of co-evolution. Table indicates average P@L for [left] predictions on native MSA; [middle] predictions after shuffling within columns (ablating co-evolution); and [right]: predictions after shuffling column order.

|  | Default | Shuffled Covariance | Shuffled Positions |
| --- | --- | --- | --- |
| MSA Pairformer | $0.71 \pm 0.21$ | $0.12 \pm 0.10$ | $0.25 \pm 0.12$ |
| MSA Transformer | $0.57 \pm 0.17$ | $0.34 \pm 0.18$ | $0.28 \pm 0.11$ |

**Figure 8:**
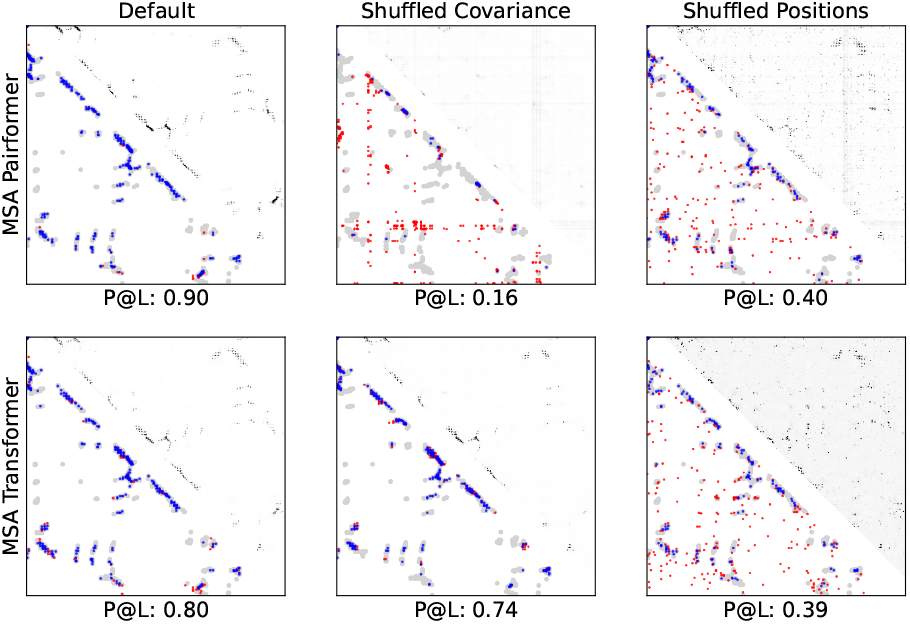
MSA Pairformer does not hallucinate contacts after removing covariance from MSAs. We compare MSA Pairformer and MSA Transformer for two types of MSA perturbations. Predicted contact maps are shown for 4MPT from the trRosetta training set. For all plots, grey indicates ground truth contacts from the crystal structure (C*β*-C*β ≤* 8Å), blue points indicate true positive predictions, and red points indicate false positive predictions for the top-L predicted contacts. [Left] shows predictions using the native MSA, [middle] shows predictions after shuffling within columns (ablating co-evolution), and [right] shows predictions after shuffling the order of the columns (maintaining co-evolution but permuting sequence order). Unlike MSA Transformer, MSA Pairformer does not hallucinate contacts after ablating co-evolution. For both models, contact accuracy mostly deteriorates when column order is shuffled.

Interestingly, whereas MSA Transformer often maintains high contact accuracy after ablating covariance (average P@L decreases from 0.57 to 0.34), MSA Pairformer does not predict contacts of the original structure (average P@L decreases from 0.71 to 0.12). This highlights MSA Pairformer’s ability to properly extract co-evolutionary signals across sequences in the MSA rather than hallucinating contacts. Additional analyses are needed to investigate the mechanisms underlying MSA Transformer’s hallucinatory behaviors.

## 5 Discussion

MSA Pairformer demonstrates that parameter efficiency and biological insight can synergistically advance protein language modeling. With just 111M parameters—less than 1% the size of ESM2-15B—the model achieves state-of-the-art performance across various biologically important tasks while uniquely capturing subfamily-specific evolutionary signals that existing MSA models miss. MSA Pairformer demonstrates substantial improvements in predicting contacts at the interface of protein-protein interactions (PPIs) and can better discriminate between binding and non-binding PPI interface sequences. Moreover, while scaling single-sequence pLMs has shown a trade-off between contact accuracy and zero-shot variant effect prediction beyond moderate model sizes, MSA Pairformer excels at both tasks.

While single-sequence pLMs have been favored for streamlining protein language modeling by circumventing MSA generation, this advantage has diminished as MSAs can now be constructed in milliseconds [58]. As MSA generation methods continue to improve, models that efficiently leverage the rapidly growing set of available sequences, and thus richer evolutionary context, are well-positioned to advance protein language modeling toward a more sustainable future. MSA Pairformer’s memory efficiency enables processing of deeper MSAs, which we find improves prediction of protein-protein interactions and variant effects (Figure S2). Crucially, unlike large single-sequence models that require expensive retraining to incorporate newly available data, MSA Pairformer can leverage such sequences directly at inference time, positioning it ideally for an era of rapidly expanding sequence databases. These results challenge the prevailing scaling paradigm and point toward a more sustainable future for protein language modeling.

MSA Pairformer’s improved performance also motivates further investigation into its underlying mechanisms. Our ablation studies reveal that triangle updates play a crucial role in disentangling direct and indirect correlations between residues. This finding provides a mechanistic explanation for MSA Pairformer’s superior contact prediction and suggests that explicitly modeling triplet relationships is essential for accurately extracting evolutionary signals from protein sequences. Furthermore, through MSA perturbation analyses, we find that unlike MSA Transformer, MSA Pairformer does not hallucinate contacts when covariance between columns of the MSA is ablated. Interestingly, Rao et al. [27] found that MSA Transformer’s ability to recover contacts after ablating covariance is directly correlated with the depth of the unfiltered MSA. In addition, we find that MSA Transformer suppresses contact predictions for strongly co-varying residues at the interface of protein-protein interactions. Together, these results suggest that MSA Transformer, akin to a single-sequence model, may sometimes retrieve co-evolutionary information stored in its parameters during training, potentially using site-wise profiles to retrieve pairwise relationships in the absence of covariance in the MSA. These results thus hint at fundamental differences in the ways by which MSA Transformer and MSA Pairformer extract evolutionary signals and motivate further investigation to comprehensively elucidate the mechanisms underlying these differences.

The absence of hallucinations in MSA Pairformer and its ability to accurately model protein-protein interactions (PPIs) opens promising directions for biological discovery. In the appendix, we show that MSA Pairformer discriminates between properly and improperly paired sequences through the presence or absence of hetero-oligomeric interface contacts, respectively (Figure S1). This result differs starkly from those uncovered by Guan and Keating [59], which demonstrated that AlphaFold3 hallucinates interactions when input MSA pairings are scrambled. This pathology may be due to the fact that AlphaFold3 was trained using erroneously paired sequences and learned to be insensitive to pairings. While this provides correct structures for examples with known interactions, it leaves AlphaFold3 susceptible to hallucinating interactions between paralogs that do not interact in reality. Although AlphaFold2/3 have been used to perform large-scale screens of PPIs, this hallucinatory behavior potentially undermines its reliability Evans et al. [60], Todor et al. [61]. MSA Pairformer is therefore better suited for large-scale screening of unknown PPIs since it models interactions directly from evolutionarily-derived statistical relationships. Furthermore, since MSA Pairformer’s pseudolikelihood scores more accurately discriminate between binding and non-binding pairs of proteins, it may improve methods that use pseudolikelihoods to pair interacting sequences in MSAs to improve PPI modeling [62].

MSA Pairformer’s ability to identify subfamily-specific signals enables exciting directions to disentangle disparate co-evolutionary signals within protein families. For example, its ability to separate subfamily sequences through bimodal attention weights in the query-biased outer product layers presents opportunities to systematically discover protein subfamilies and their distinct functional properties. The query-biased outer product may also enhance supervised structure prediction models like AlphaFold3 by enabling characterization of subfamily-specific structural differences that current approaches average out. Interestingly, recent studies have demonstrated that sub-sampling MSAs via clustering can facilitate exploration of alternative conformations [33, 34]. By learning the evolutionary relationships between sequences within a protein family, MSA Pairformer may enable further exploration of conformational landscapes encoded in distinct co-evolutionary patterns in MSAs.

The ProteinGym results reveal an intriguing trend in how different protein language model architectures scale. While single-sequence pLMs improve at contact prediction as they grow larger, they struggle to maintain variant effect prediction performance. In contrast, MSA-based models like MSA Pairformer and MSA Transformer achieve strong performance across both tasks. These results may reflect a fundamental difference in how single-sequence and MSA models encode evolutionary information. As single-sequence models scale, they appear to narrow their focus to specific protein family clusters, improving sequence recovery, perplexity, and contact prediction but losing the global evolutionary statistics, which may be crucial for variant effect prediction [63]. Conversely, MSA Pairformer’s smooth sequence weight distributions suggest it retains a more global view of evolutionary signals while still biasing extraction toward query-relevant information. This balanced approach may explain its strong performance across diverse tasks. Further investigation is needed to determine whether this architectural advantage persists as MSA models continue to improve.

Overall, MSA Pairformer demonstrates that parameter-efficient models can achieve state-of-the-art performance and open avenues for biological discovery. By learning to focus on query-relevant evolutionary signal, the model opens new avenues for both computational efficiency and biological discovery in the era of rapidly expanding genomic databases.

## 6 Limitations

Given the vast range of biological questions that may be approached using pLM embeddings, we are eager to find how MSA Pairformer can support advances in protein modeling and design beyond the scope of this study. Moreover, while our ablation studies suggest a potential role of triangle updates in MSA Pairformer, additional analyses are needed to comprehensively investigate MSA Pairformer’s internal mechanisms and how they contribute to the model’s broad performance gains. Furthermore, while this work introduces a variant of softmax attention, pre-softmax differential attention, we do not fully explore its properties or limitations. Future work can further investigate its behavior across domains to better understand its strengths and potential applications.

## 7 Acknowledgments

The authors thank Chenxi Ou, Claudia Chu, Eric Guan, James Roney and Olivia Tang and the entire Ovchinnikov lab for valuable discussions, feedback, and support throughout the project. We also thank Roshan Rao for providing the MSA Transformer and ESM2 contact head training data. Y.A. was funded by the Schimmel Family Program for Life Sciences, Z.Z. acknowledges the Swiss National Science Foundation Postdoctoral Mobility grant (P500PB_225547), and S.O. acknowledges funding from NSF grant MCB2032259 and Amgen. Y.A. and S.O. were also supported by MIT Lincoln Laboratory Advanced Concepts Committee. M.S. acknowledges support by the National Research Foundation of Korea (grant RS-2024-00396026), and Novo Nordisk Foundation (NNF24SA0092560). M.M. acknowledges support from the National Research Foundation of Korea (grant RS-2023-00250470).

## 8 Code availability

Code and model weights are available at https://github.com/yoakiyama/MSA_Pairformer and https://huggingface.co/yoakiyama/MSA-Pairformer.

A Google Colaboratory notebook for exploratory use is available at https://colab.research.google.com/github/yoakiyama/MSA_Pairformer/blob/main/MSA_Pairformer_with_MMseqs2.ipynb. The notebook provides automated MSA generation using an updated version of the ColabFold MMseqs2 protocol, which we detail in the appendix.

## Notes

https://github.com/yoakiyama/MSA_Pairformer

### Competing Interest Statement

The authors have declared no competing interest.

