## Appendix for "Scaling down protein language modeling with MSA Pairformer"

### A Technical Appendices and Supplementary Material

#### A.1 Building paired MSAs using MMseqs2

For the protein complex interface contact prediction analyses, we used the same MSAs as Ovchinnikov et al. [1]. To enable broader accessibility and automated MSA generation, we also provide support for pairing MSAs using MMSeqs2 Steinegger and Söding [2], Mirdita et al. [3, 4], Kallenborn et al. [5], Mirdita et al. [6], Steinegger and Söding [7]. By default, ColabFold pairs proteins according to the highest MMseqs2 alignment score per taxonomic identifier, similar to the protocol used in AlphaFold-Multimer [6, 8]. To improve orthology inference, we introduce a proximity-based pairing scheme that infers genomic proximity directly from protein accessions [1]. First, we translate UniProt and UniParc accessions into structured integers, based on the convention that sequentially numbered accessions reflect neighboring genes [9–13]. Beginning from the highest scoring protein identified through an MMseqs2 search against the UniRef100, we iteratively select the closest unmatched protein within a predefined numerical threshold (default distance $\leq 20$). This process is repeated until all suitable protein pairs have been identified. We allow either greedily pairing all possible matches or enforcing that all protein chains must be covered by paired database proteins. Additionally, we improve the sensitivity of protein pairing by incorporating a profile-to-sequence alignments computed with MMseqs2. For example, for the dimer 1TYG, the number of pairable proteins increased from approximately one hundred to over two thousand pairs.

To demonstrate the importance and efficacy of proximity-based pairing, we highlight the ABC transporter ModBC in complex with its binding protein ModA (PDB ID: 2ONK) (Figure S1). The ABC transporter family contains many paralogous genes per species, which form different hetero-oligomeric complexes [14]. Therefore, mispairing sequences may prevent aberrant modeling of the hetero-oligomeric interaction. Using 512 sequences, we predict contacts using the default ColabFold pairing method, the default method with coverage and minimum query identity filtering (coverage $\geq 75\%$ and minimum sequence identity with query $\geq 30\%$), and MMseqs2 proximity-based pairing with coverage and query identity filtering (same as previous). We find that hetero-oligomeric interactions are only captured after pairing sequences, demonstrating the importance of properly pairing sequences to model PPIs. Furthermore, these results highlights MSA Pairformer's ability to discriminate between properly and improperly paired sequences through interfacial contacts, as it does not hallucinate contacts when sequences are mispaired.

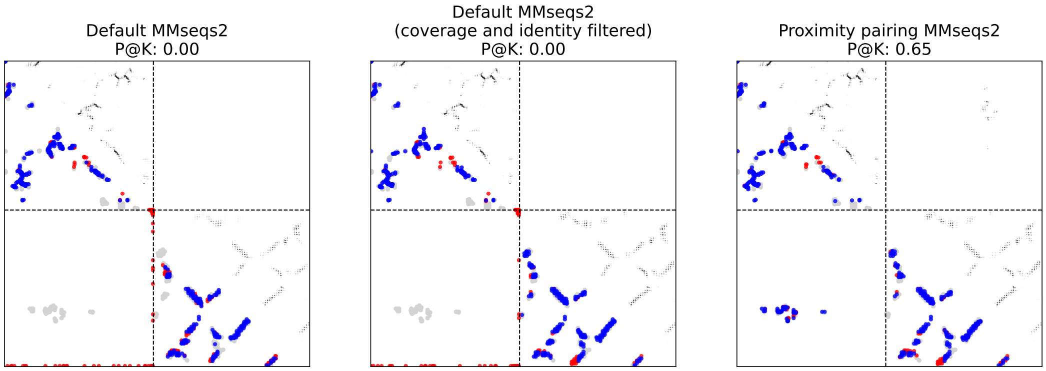


Figure S1: Pairing sequences using genomic proximity is critical for modeling protein-protein interactions.

#### A.2 Impact of MSA depth on model performance across tasks

We investigate the effect of MSA depth on unsupervised contact, PPI, and variant effect prediction.

Using a held out set of 100 targets from the trRosetta training set, we increased MSA depth from 1 to 512, incrementing by powers of 2. We find that while the average P@L increases monotonically, it increases sharply up to 0.59 with 32 sequences and begins to saturate at roughly 0.72 using 128 sequences. We similarly test the effects of MSA depth on PPI prediction using the same 31 interactions from Ovchinnikov et al. [1]. We find that mean interface contact precision continues to increase up to 0.48 using 256 sequences, at which point performance begins to plateau. These results suggest that more sequences are needed in order to accurately predict hetero-oligomeric interactions.

Lastly, we investigate the effects of MSA depth on variant effect prediction. We compute the average Spearman's correlation on the 217 ProteinGym deep mutational scan experiments using 400, 2048, and 4096 sequences, respectively. We find that performance monotonically increased for these three MSA depths, suggesting that comprehensively incorporating evolutionary context may be particularly important for predicting variant effects.

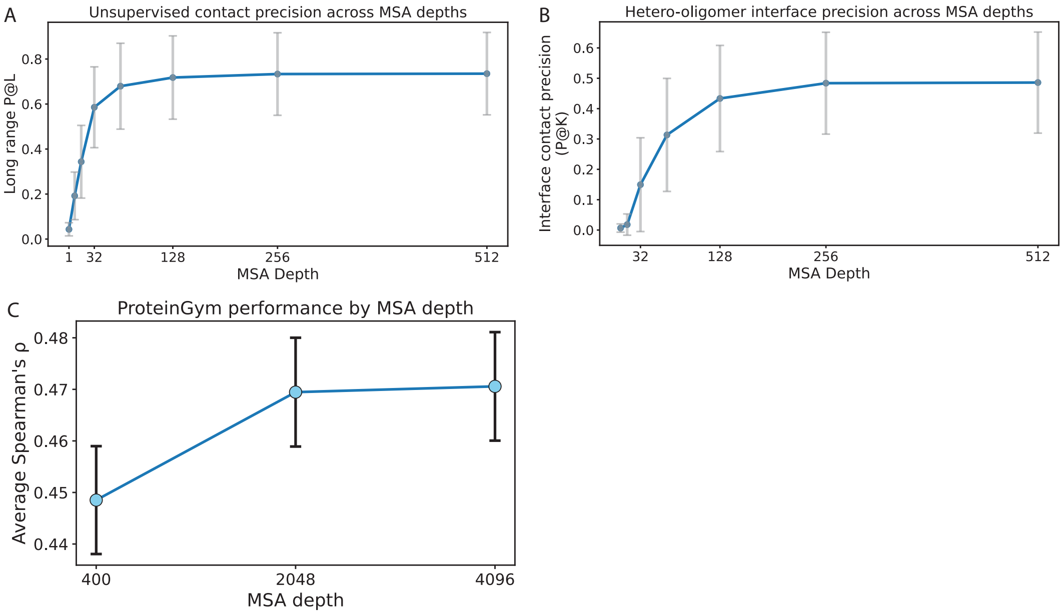


Figure S2: The effects of MSA depth on performance varies across tasks. We evaluate the effects of varying MSA depth across three tasks: A) unsupervised contact prediction, B) protein-protein interaction prediction, and C) variant effect prediction.

#### A.3 Time and memory complexity

MSA Pairformer demonstrates superior memory efficiency compared to MSA Transformer across both MSA depth and sequence length dimensions, using $O(SL + L^2)$ memory compared to $O(SL^2 + S^2L)$, where $S$ is the MSA depth and $L$ is the sequence length. To evaluate their capacity for processing deeper MSAs—which provide broader evolutionary context—we generated MSAs with 1024 residues (MSA Transformer's maximum length) and varied the number of sequences by powers of two. While MSA Transformer exhausts available GPU memory (96GB) at 384 sequences, MSA Pairformer successfully processes 4096 sequences using only 74GB—over a 10-fold increase in capacity (Figure **??**A). We conducted a complementary experiment varying sequence length for MSAs containing 1024 sequences (MSA Transformer's maximum depth). MSA Transformer reaches its memory limit at just 128 residues, whereas MSA Pairformer handles sequences up to 2304 residues using only 60GB (Figure **??**B). These results demonstrate that MSA Pairformer achieves more than an order of magnitude improvement in memory efficiency, enabling the processing of substantially deeper and longer MSAs crucial for capturing comprehensive evolutionary information.

MSA Pairformer runs in $O(SL^2 + L^3)$ versus MSA Transformer's $O(SL^2 + S^2L)$. To demonstrate this complexity in practice, we conduct similar experiments as before. To measure the effects of sequence length, we use a fixed MSA depth of 384 sequences and vary the sequence length up to 1024 residues S3

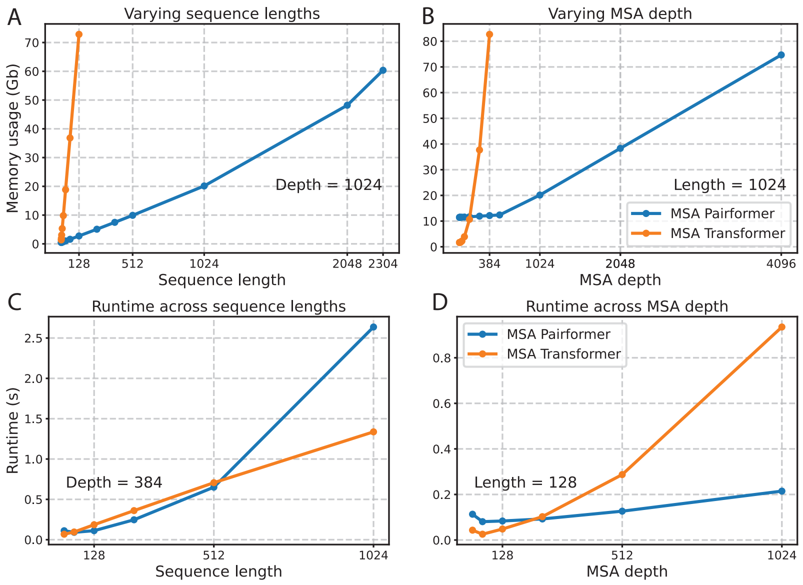


Figure S3: MSA Pairformer achieves efficiency gains over MSA Transformer. Memory usage of MSA Pairformer and MSA Transformer across varying (A) sequence lengths for a 1024 sequence MSA; and (B) MSA depths for a 1024 residue sequence. Runtime of MSA Pairformer and MSA Transformer across varying (C) sequence lengths for a 384 sequence MSA; and (D) MSA depths for a 128 residue sequence. All experiments were run on an NVIDIA H100 GPU.

##### A.4 Cropping relative positional embeddings

In order to better understand why MSA Pairformer outperforms models like MSA Transformer in contact prediction, we investigated the potential role of positional embeddings. Whereas MSA Transformer uses learned absolute positional embeddings, MSA Pairformer uses learned relative positional embeddings, cropped to a maximum distance of 32 residues. To investigate whether this cropping meaningfully contributes to the model's performance, we

1. Fine-tuned MSA Pairformer for an additional 5000 steps.

2. Fine-tuned MSA Pairformer for an additional 5000 steps, expanding the learned relative positional embedding from 32 to 320, which is the maximum training size. For this model, we initialize any distance greater than 32 to the projection for 32 residue separation.

We find that the training, validation, and test losses are approximately identical, and the long-range contact P@L on a held out set of 1000 proteins from the trRosetta training set were also approximately identical. These results suggest that extending the maximum relative distances may not have much of a consequence. For MSA Pairformer, cropping these relative distances to a maximum value of 32 residue separation offers simplicity and reduces the complexity of the positional embedding. We note that training the model from scratch with different cropping values may lead to more separation in the models, but we leave this analysis for future work.

##### A.5 Training details

###### A.5.1 Data

We use the filtered set of 270k alignments from the OpenFold OpenProteinSet. These alignments were generated by searching every cluster in Uniclust30 (v. 2018-08). Additional details of the dataset can be found in the OpenProteinSet and OpenFold manuscripts [15, 16].

##### A.5.2 Base model pre-training

We first pre-train a base model that uses uniform sequence weighting in the outer product, as in the outer product mean of AlphaFold2/3. This provides us a baseline and starting point for testing different attention mechanisms for the query-biased outer product layers.

The model takes as input a batch of partially masked and mutated MSAs and is trained to recover masked/mutated tokens. Following the training procedures of ESM2 and MSA Transformer, 15% of tokens are selected, among which 80% are masked, 10% are mutated to a random amino acid, and 10% remain unaltered.

We train with an effective batch size of 12 using gradient accumulation for 50,000 steps. We use the AdamW optimizer with parameters $\beta_1 = 0.95$, $\beta_2 = 0.99$, $\epsilon = 10^{-8}$, and $\lambda = 0.1$. The base learning rate is $8 * 10^{-4}$ and is linearly increased from 0 for 1,000 steps, then linearly decreased to $8 * 10^{-5}$ for the remaining gradient updates. We apply gradient clipping if the global norm exceeds 1.

##### A.5.3 Query-biased attention fine-tuning

After pre-training the base model using uniform sequence weighting, we fine-tuned the model to compute a weighted average outer product based on each sequence's relevance to the query using normalized attention weights. We refer to this as the query-biased outer product (Figure 1A). This approach specifically addresses the key limitation in current MSA models: the averaging of co-evolutionary information that can obscure subfamily-specific signals critical for understanding protein function and structure. During fine-tuning, we used a modified masking strategy where only the query sequence (first row of the MSA) was partially masked, while homologous sequences remained fully visible to provide evolutionary context for learning query-biased attention weights.

We experiment with four methods to compute attention: 1) standard scaled-dot product attention, 2) differential attention, 3) non-negative differential attention and 4) pre-softmax differential attention. We describe these methods in the following sections.

Recall the MSA representation $x \in \mathbb{R}^{M \times L \times d_m}$, where $M$ represents the number of sequences, $L$ the sequence length, and $d_m$ the embedding dimension. For each token representation $x_{s,i}$, we project it to a key vector $k_{s,i} \in \mathbb{R}^{''}$. For each token representation in the query sequence, we project it to a query vector $q_{0,i}$. We compute the dot products between the query and key vectors across the same column, then mean-pool the dot products to obtain a single, $M$-dimensional attention weight vector. We denote the mean-pooled dot products $qK^T \in \mathbb{R}^M$.

$$qK^T = \frac{\sum_{i=1}^{L} q_{0,i} K_i^T}{L} \tag{1}$$

For differential and pre-softmax differential attention, we compute two of these vectors for reasons explained in the following sections.

**Differential attention** To improve sequence weighting accuracy, we explored alternatives to standard dot-product attention [17]. Introduced by Ye et al. [18] via the Differential Transformer, differential attention aims to remove noise from attention weights. Whereas standard scaled dot-product attention computes a single attention matrix per attention head, differential attention computes two attention matrices, computing the final attention matrix as the difference between the two. The first softmax attention matrix is analogous to the attention matrix from standard attention, and it may apply non-trivial attention to irrelevant context due to noise. The second softmax attention matrix can be thought of as removing this noise. The difference between standard dot-product attention and differential attention is shown below.

$$\text{Attn}(X) = \text{softmax}(\frac{qK^T}{\sqrt{d_q}}) \tag{2}$$

$$\text{DiffAttn}(X) = \text{softmax}(\frac{q_1 K_1^T}{\sqrt{d_q}}) - \lambda * \text{softmax}(\frac{q_2 K_2^T}{\sqrt{d_q}}) \tag{3}$$

$\lambda$ is a learned scaling factor and is reparameterized as

$$\lambda = \exp(\lambda_{q_1} \cdot \lambda_{k_1}) - \exp(\lambda_{q_2} \cdot \lambda_{k_2}) + \lambda_{\text{init}}$$

where $\lambda_{q_1}, \lambda_{k_1}, \lambda_{q_2}, \lambda_{k_2} \in \mathbb{R}^d$ are learnable vectors. $\lambda_{\text{init}} \in (0, 1)$ is a constant used to initialize $\lambda$ and is set via $\lambda_{\text{init}} = 0.8 - 0.6 \times \exp(-0.3 \cdot (l - 1))$, where $l \in [1, N]$ represents the layer index. Conceptually, this scaling factor enables the model to adapt the level of noise cancellation for each layer.

**Pre-softmax differential attention** While the Differential Transformer showed improvements over standard transformer attention in language modeling, we hypothesize that a slight modification may provide additional benefits.

We note a few potential limitations of differential attention. By computing the differential after applying softmax to both dot product matrices, differential attention does not rescue diminished attention for relevant tokens caused by attention weights assigned to irrelevant tokens in the first attention matrix. To illustrate this point, suppose we have a sequence of tokens $x_i$ for $i = 0, ..., N$, and 100% of the attention should be applied to token $x_0$. Next, suppose that after applying softmax to the first dot product matrix, the sum of the normalized attention weights assigned to tokens $x_1, ..., x_N$ is 0.1. While differential attention may reduce the weights assigned to tokens $1, \cdots, N$ to 0, the final attention weight applied to $x_0$ is at most $\text{attn}_0 \leq 0.9$. This pathology is related to the attention dispersion problem described by Veličković et al. [19].

Furthermore, Differential Attention may result in negative attention weights. In the context of weighting sequences in an MSA, negative values may result in unintuitive sequence weightings and instability.

To circumvent this down-weighting effect and to prevent negative attention weights, we explore a simple modification, computing the difference before applying softmax.

$$\text{PresoftmaxDiffAttn}(X) = \text{softmax}(\frac{q_1 K_1^T - \lambda q_2 K_2^T}{\sqrt{d_q}}) \tag{4}$$

This formulation is similar to Selective Attention, introduced by Leviathan et al. [20], where an auxiliary attention matrix $F$ is computed as a cumulative sum of masking weights derived from an attention matrix. Using Selective Attention in an autoregressive transformer model allows tokens to mask previous tokens that are considered unneeded for future tokens, thereby reducing the context window and preventing performance degradations attributed to noise. In Selective Attention, $F$ is constrained to be non-negative such that this matrix can only mask, not upweight, tokens. The pre-softmax differential used in this work can thus be thought of as a hybrid between differential and selective attention. Here, we do not apply non-negativity constraints to either dot product matrix, allowing both $\lambda$ and $q_2 K_2^T$ to take negative and positive values.

We also note that this pre-softmax differential can be rewritten as

$$\begin{aligned}
\text{PresoftmaxDiffAttn}(X) &= \text{softmax}\left(\frac{(X_0 W_{Q_1})(X W_{K_1})^T - \lambda (X_0 W_{Q_2})(X W_{K_2})^T}{\sqrt{d_q}}\right) &(5)\\
&= \text{softmax}\left(\frac{X_0 W_{Q_1} W_{K_1}^T X^T - \lambda X_0 W_{Q_2} W_{K_2}^T X^T}{\sqrt{d_q}}\right) &(6)\\
&= \text{softmax}\left(\frac{X_0 (W_{Q_1} W_{K_1}^T - \lambda W_{Q_2} W_{K_2}^T) X^T}{\sqrt{d_q}}\right) &(7)
\end{aligned}$$

This decomposition reveals that pre-softmax differential attention is mathematically equivalent to standard attention with an effective projection matrix $W_Q W_K^T = (W_{Q_1} W_{K_1}^T - \lambda W_{Q_2} W_{K_2}^T)$. However, the learnable scaling factor $\lambda$ may provide additional flexibility in controlling attention sharpness that is otherwise constrained by layer normalization and weight decay, as described by Veličković et al. [19]. Therefore, this formulation may effectively enable tuning the attention sharpness while preserving the stability and generalization benefits of QK-Norm and weight decay [19, 21–23].

We test this pre-softmax differential against the other attention variants in the following section.

**Experiments using the attention variants in the query-biased outer product** We fine-tune the pre-trained base model using the four attention variants:

- Standard scaled dot product attention
- Differential attention
- Non-negative differential attention
  - A simple modification of differential attention. We apply ReLU to the final attention vector and then re-normalize so that the weights sum to 1
- Pre-softmax differential attention

In order to isolate the effect of the attention mechanisms, we standardize the experimental conditions by using identical random seeds, optimization parameters, and training procedure for each method. Specifically, we trained 12,500 gradient steps, using the AdamW optimizer with parameters $\beta_1 = 0.9, \beta_2 = 0.999$, and $\lambda = 0.001$. We use a maximum learning rate of $5 * 10^{-4}$ and $5 * 10^{-5}$ for attention related parameters and all other parameters, respectively. Learning rate is increased for 500 steps and decayed by a factor of $10^5$ for 12,000 steps using a cosine learning rate scheduler.

We evaluated performance using validation perplexity during fine-tuning (Figure S4). All three differential attention variants outperformed standard scaled dot-product attention, with pre-softmax differential achieving the lowest validation perplexity, followed by non-negative differential attention, then differential attention. The performance ranking (pre-softmax > non-negative differential > differential > standard) suggests that both preventing negative weights and computing differences before normalization contribute to improved sequence weighting accuracy.

The final version of MSA Pairformer used for downstream analyses uses pre-softmax differential attention and was trained for an additional 5,500 steps steps using the same hyperparameters. As demonstrated in Section 4, query-biased outer product translates to gains in both contact prediction accuracy and the ability to extract subfamily-specific co-evolutionary signals that are critical for capturing relevant context when building sequence and pairwise representations.

In this work, we focus on protein language modeling and leave further analyses comparing the advantages, limitations, and performance differences between these attention variants to future work.

**Algorithm 1** Query-biased outer product

**def** QuerybiasedOuterProduct($\{\mathbf{m}_{si}\}$, $c = 16$, $c_z = 256$):
$\mathbf{m}_{si} \leftarrow$ LayerNorm($\mathbf{m}_{si}$)
$\mathbf{b}_{si}, \mathbf{c}_{si} =$ LinearNoBias($\mathbf{m}_{si}$) ▷ $\mathbf{a}_{si}, \mathbf{b}_{si} \in \mathbb{R}^c$
$\mathbf{q}_{1i}, \mathbf{q}_{2i} =$ LinearNoBias($\mathbf{m}_{si}$) ▷ $\mathbf{q}_{1i}, \mathbf{q}_{2i} \in \mathrm{R}^{d_q}$
$\mathbf{K}_{1si}, \mathbf{K}_{2si} =$ LinearNoBias($\mathbf{m}_{si}$) ▷ $\mathbf{K}_{1si}, \mathbf{K}_{2si} \in \mathrm{R}^{d_q}$
$\mathbf{q}_{1i}, \mathbf{q}_{2i} \leftarrow$ LayerNorm(concat($\mathbf{q}_{1i}, \mathbf{q}_{2i}$))
$\mathbf{K}_{1si}, \mathbf{K}_{2si} \leftarrow$ LayerNorm(concat($\mathbf{K}_{1si}, \mathbf{K}_{2si}$))
$\mathbf{h_{1si}}, \mathbf{h_{2si}} = \mathbf{q_{1i}} \cdot \mathbf{K_{1si}}, \mathbf{q_{2i}} \cdot \mathbf{K_{2si}}$
$\mathbf{h}_{1s}, \mathbf{h}_{2s} \leftarrow \frac{1}{n}\sum_{i=1}^{n} \mathbf{h}_{1si}, \frac{1}{n}\sum_{i=1}^{n} \mathbf{h}_{2si}$
$\lambda = \exp(\lambda_{\mathbf{q}_1} \cdot \lambda_{\mathbf{k}_1}) - \exp(\lambda_{\mathbf{q}_2} - \lambda_{\mathbf{k}_2}) + \lambda_{\text{init}}$
$\mathbf{h}_s = \mathbf{h}_{s1} - \lambda \mathbf{h}_{s2}$
$\mathbf{a}_s =$ softmax($\mathbf{h}_s$)
$\mathbf{o}_{ij} =$ flatten($\sum_s \mathbf{a}_s * \mathbf{b}_{si} \otimes \mathbf{c}_{sj}$) ▷ $\mathbf{o}_{ij} \in \mathrm{R}^{c*c}$
$\mathbf{s}_{ij}, \mathbf{g}_{ij} =$ Linear($\mathbf{o}_{ij}$) ▷ $\mathbf{s}_{ij}, \mathbf{g}_{ij} \in \mathrm{R}^{c_z}$
$\mathbf{z}_{ij} \leftarrow$ LinearNoBias(swish($s_{ij}$) $\odot$ $g_{ij}$) ▷ $\mathbf{z}_{ij} \in \mathrm{R}^{c_z}$
**return** $\mathbf{z}_{ij}$

##### A.6 Fitting the contact head for unsupervised contact prediction

Previous studies have shown that attention maps from protein language models can be used to predict residue contacts [24, 25]. Following a similar procedure as [24], we fit a small contact head using a training set of 20 targets from the trRosetta training set. We define contacts as residue pairs with C$\beta$-C$\beta$ < 8Å. For glycines, the C$\beta$ is placed based on the N, CA, C vector of the backbone, following the protocol of TrRosetta [26]. Here, we deviate slightly from MSA Transformer and ESM2 by fitting

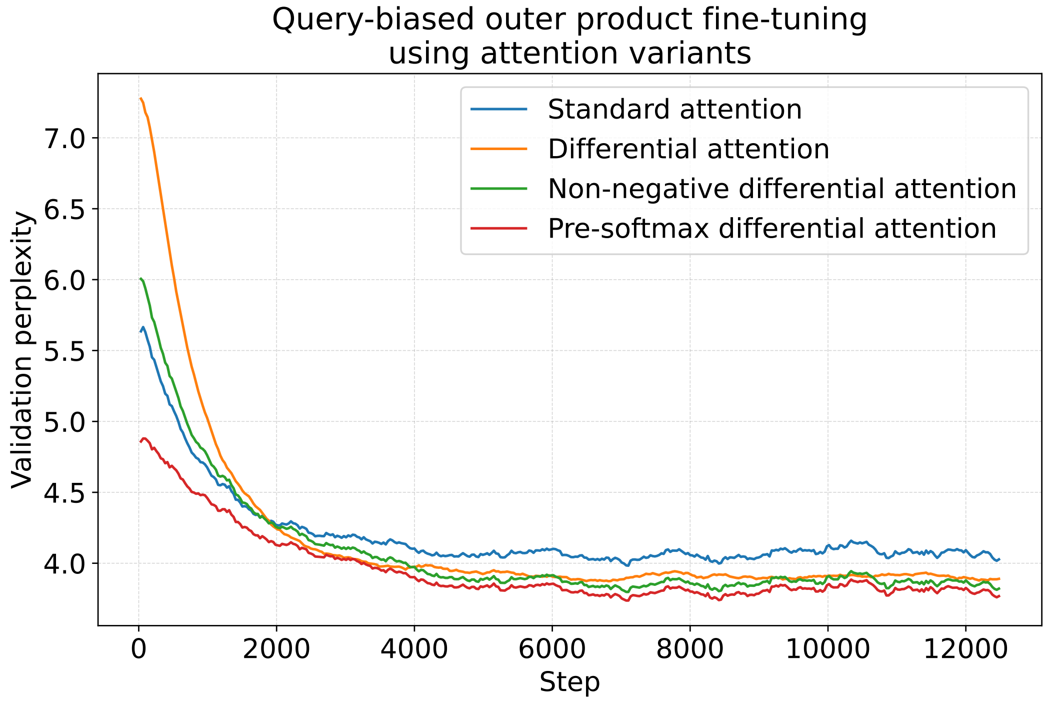


Figure S4: Validation perplexity across training steps for query-biased outer product fine-tuning using different attention variants.

the contact head on the pair representation after each layer in the model rather than stacking the attention weights across the model.

---

**Algorithm 2** Contact head

---

$\mathbf{z}_{ij} \leftarrow \text{LayerNorm}(\mathbf{z}_{ij})$ ▷ $\mathbf{z}_{ij} \in \mathbb{R}^{d_p}$

$\mathbf{c}_{ij} \leftarrow \text{sigmoid}(\text{Linear}(\mathbf{z}_{ij}))$ ▷ $\mathbf{c}_{ij} \in \mathbb{R}$

$\mathbf{c}_{ij} \leftarrow \frac{\mathbf{c}_{ij}+\mathbf{c}_{ji}}{2}$

**return** $\mathbf{c}_{ij}$

---

We find that contact precision maximizes between layers 12 and 16. For all evaluations, we use the pair representation at layer 15. For the final contact prediction head, we performed a hyperparameter sweep of learning rate and weight decay, ultimately using $\eta = 0.01$ and $\lambda = 0.13$.

##### A.7 CASP15 unsupervised contact prediction

We evaluate unsupervised contact prediction using the 49 monomeric targets from CASP15. In order to include MSA Transformer in the evaluation, we limited the analysis in Figure 2A to 46 sequences with fewer than 1024 residues (T1154, T1158, and T1169 are removed at this step). We generate MSAs for the targets by searching Uniref30 (ver. Feb 2023) using hhblits. We use an e-value cutoff of 0.001 and four iterations. In order to control for the composition of the MSAs provided to MSA Pairformer and MSA Transformer, we limit the the MSAs to a maximum depth of 512 sequences. We use hhfilter, removing sequences with less than 50% coverage, less than 15% sequence identity to the query, and setting a maximum pairwise sequence identity of 95%, a minimum. If hhfilter returns an MSA with more than 512 sequences, we apply the greedy diversity maximization approach used by Rao et al. [24] to subset to 512 sequences.

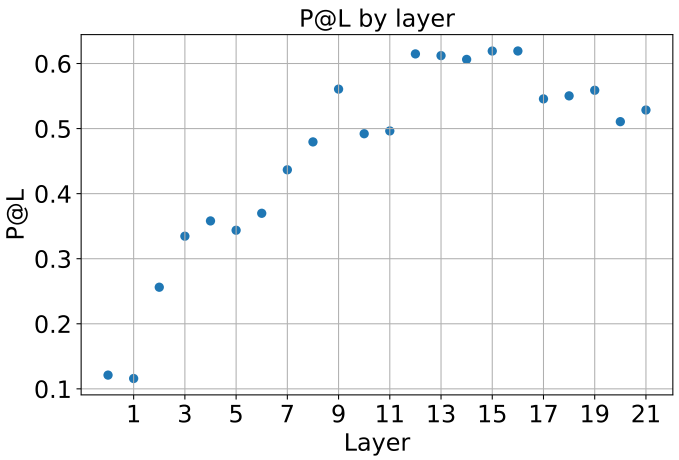


Figure S5: Validation long-range P@L across layers of MSA Pairformer. Final model uses the pair representation at layer 15 to predict contacts.

Table S1: CASP15 long-range P@L across protein structure prediction methods

| | Long-range P@L |
|---|---|
| MSA Pairformer | **0.52** |
| MSA Pairformer (uniform sequence weights) | 0.50 |
| MSA Transformer | 0.44 |
| ESM2 15B | 0.46 |
| ESM2 3B | 0.45 |
| ESM2 650M | 0.42 |
| ESM2 150M | 0.34 |

##### A.8 Response regulator domain analysis

Following Malinverni and Barducci [27], generated the response regulator (RR) domain MSA by first downloading all aligned sequences for the RR family (PFAM ID: PF00072), as well as the aligned sequences for the DNA binding domains of the three subfamilies, OmpR (PFAM ID: PF00486), LytTR (PFAM ID: PF04397) and GerE (PFAM ID: PF00196). We then subset the RR family alignment by selecting sequences that are included in one of the three subfamily alignments. Lastly, we aligned the canonical sequences used in the crystallographic structures (PDB IDs: 1NXS, 4CBV and 4E7P for OmpR, LytTR and GerE, respectively) to the existing alignment in order to include the sequences.

In order to evaluate whether MSA Pairformer extracts subfamily-specific co-evolutionary signals, we subsetted the MSA for 4096 sequences using hhfilter. For each subfamily prediction, we assign the PDB canonical sequence as the query and predict contacts. Noteably, using uniform sequence weighting results in approximately the same contact predictions regardless of the query sequence, whereas query-biased outer product allows for better prediction of the appropriate interface contacts. In TableS2, TableS3 and TableS4, we show the top ten predicted interface contacts using MSAs with the OmpR, GerE, and LytTR sequences as the query sequence, respectively. LytTR contacts are not fully recovered, though non-LytTR inter-chain contacts generally show decreased probability compared to uniform weighted averaging. A previous study also found that pLMs struggle to extract these contacts [28].

We repeat this analysis using MSA Transformer. Importantly, its tied row attention averages the attention maps, and therefore its inferred pairwise dependencies, across the sequences in an MSA. We therefore do not expect MSA Transformer to isolate subfamily-specific signals relevant to the query sequence. We sample an MSA of 1024 sequences containing 611, 312, and 100 sequences from GeRE, LytTR, and OmpR, respectively, using hhfilter and diversity maximization. We assign a sequence from each subfamily and predict contacts. Indeed, MSA Transformer's contact predictions do not accurately reflect changes in the query sequence (Figure S6). For example, regardless of which subfamily the query sequence belongs to, MSA Transformer predicts the same 3 GerE contacts. Interestingly, for the OmpR query sequence, MSA Transformer only predicts a single OmpR contact, despite it predicting 2 and 3 OmpR contacts when the query sequence belongs to the LytTR and GerE subfamilies, respectively. This result highlights MSA Transformer's complete inability to separate or prioritize the correct subfamily-specific signals.

Overall, our results therefore highlight the importance of the query-biased outer product in MSA Pairformer for capturing subfamily-specific properties.

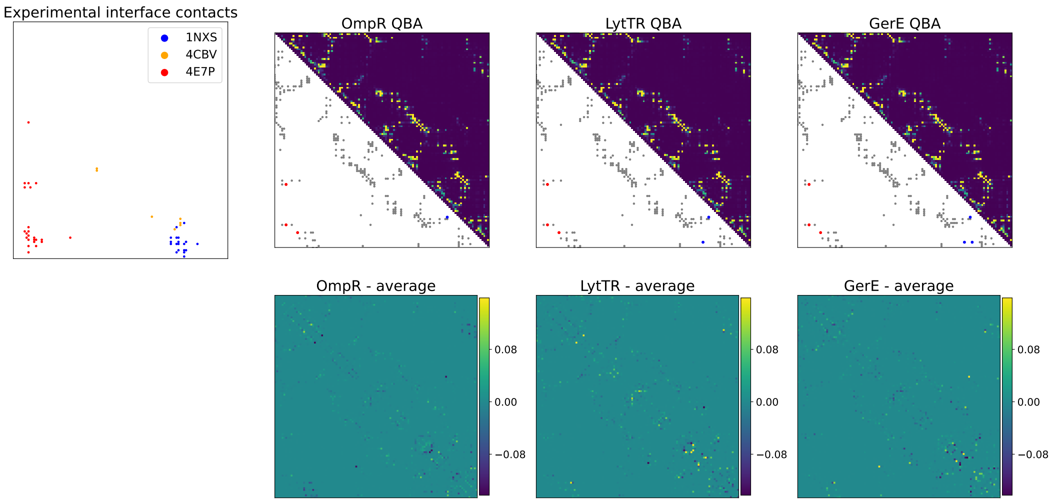


Figure S6: MSA Transformer fails to isolate subfamily-specific signals from the response regulator superfamily.

Table S2: Top ten interface predictions for the OmpR query sequence using MSA Pairformer. Contact prediction probabilities using both query-biased outer product and uniform sequence weighting are provided. 9 of the 10 top predictions belong to the correct subfamily.

| Interface contact subfamily | Interface contact | Contact prediction query-biased | Contact prediction uniform weighting |
|---|---|---|---|
| OmpR-1NXS | (90, 96) | 0.70 | 0.63 |
| OmpR-1NXS | (90, 110) | 0.59 | 0.28 |
| OmpR-1NXS | (90, 109) | 0.56 | 0.14 |
| OmpR-1NXS | (83, 106) | 0.53 | 0.02 |
| OmpR-1NXS | (86, 106) | 0.47 | 0.01 |
| OmpR-1NXS | (87, 106) | 0.45 | 0.04 |
| OmpR-1NXS | (83, 103) | 0.44 | 0.08 |
| GerE-4E7P | (6, 100) | 0.41 | 0.34 |
| OmpR-1NXS | (87, 109) | 0.36 | 0.02 |
| OmpR-1NXS | (90, 106) | 0.28 | 0.22 |

Table S3: Top 10 interface predictions for the GerE query sequence using MSA Pairformer. Contact prediction probabilities using both query-biased outer product and uniform sequence weighting are provided. Among the top 10 predicted contacts, 7 belong to the correct subfamily. For contacts that belong to a different subfamily, query-biased outer product reduces contact probabilities.

| Interface contact subfamily | Interface contact | Contact prediction query-biased | Contact prediction uniform weighting |
|---|---|---|---|
| OmpR-1NXS | (90, 96) | 0.46 | 0.63 |
| GerE-4E7P | (11, 104) | 0.46 | 0.14 |
| GerE-4E7P | (6, 100) | 0.41 | 0.34 |
| GerE-4E7P | (8, 104) | 0.36 | 0.08 |
| GerE-4E7P | (12, 104) | 0.34 | 0.30 |
| GerE-4E7P | (6, 79) | 0.24 | 0.24 |
| GerE-4E7P | (8, 100) | 0.19 | 0.07 |
| OmpR-1NXS | (90, 110) | 0.16 | 0.28 |
| GerE-4E7P | (9, 79) | 0.11 | 0.12 |
| LytTR-4CBV | (8, 102) | 0.10 | 0.14 |

Table S4: Top 10 interface predictions for the LytTR query sequence. Contact prediction probabilities using both query-biased outer product and uniform sequence weighting are provided. Among the top 10 predicted contacts, only 1 belongs to the correct subfamily. For contacts that belong to a different subfamily, query-biased outer product generally shows lower probabilities.

| Interface contact subfamily | Interface contact | Contact prediction query-biased | Contact prediction uniform weighting |
|---|---|---|---|
| OmpR-1NXS | (90, 96) | 0.52 | 0.63 |
| OmpR-1NXS | (90, 106) | 0.34 | 0.22 |
| GerE-4E7P | (6, 100) | 0.31 | 0.34 |
| GerE-4E7P | (6, 79) | 0.31 | 0.24 |
| GerE-4E7P | (12, 104) | 0.26 | 0.30 |
| GerE-4E7P | (9, 79) | 0.19 | 0.12 |
| OmpR-1NXS | (90, 110) | 0.19 | 0.28 |
| OmpR-1NXS | (90, 109) | 0.15 | 0.14 |
| OmpR-1NXS | (87, 106) | 0.11 | 0.08 |
| LytTR-4CBV | (8, 102) | 0.10 | 0.14 |

Table S5: Top ten interface predictions for the OmpR query sequence using MSA Transformer. We compare to the average contact predictions using the LytTR and GerE sequences.

| Interface contact subfamily | Interface contact | Contact prediction | Contact prediction LytTR-GeRE average |
|---|---|---|---|
| OmpR-1NXS | (90, 96) | 0.89 | 0.92 |
| GerE-4E7P | (6, 100) | 0.40 | 0.41 |
| GerE-4E7P | (6, 79) | 0.40 | 0.42 |
| GerE-4E7P | (12, 104) | 0.19 | 0.20 |
| OmpR-1NXS | (91, 109) | 0.15 | 0.14 |
| OmpR-1NXS | (87, 109) | 0.14 | 0.23 |
| OmpR-1NXS | (90, 109) | 0.11 | 0.12 |
| LytTR-4CBV | (85, 99) | 0.05 | 0.06 |
| GerE-4E7P | (15, 104) | 0.05 | 0.05 |
| GerE-4E7P | (9, 79) | 0.03 | 0.04 |

##### A.9 ProteinGym zero-shot variant effect prediction

We evaluate MSA Pairformer on zero-shot variant effect prediction using the ProteinGym DMS substitution benchmark. Using the MSAs provided by ProteinGym, we subset a maximum of 4096

Table S6: Top ten interface predictions for the LytTR query sequence using MSA Transformer. We compare to the average contact predictions using the OmpR and GerE sequences.

| Interface contact subfamily | Interface contact | Contact prediction | Contact prediction OmpR-GerE average |
|---|---|---|---|
| OmpR-1NXS | (90, 96) | 0.92 | 0.91 |
| GerE-4E7P | (6, 100) | 0.43 | 0.41 |
| GerE-4E7P | (6, 79) | 0.40 | 0.42 |
| OmpR-1NXS | (87, 109) | 0.25 | 0.17 |
| GerE-4E7P | (12, 104) | 0.21 | 0.20 |
| OmpR-1NXS | (91, 109) | 0.15 | 0.15 |
| OmpR-1NXS | (90, 109) | 0.13 | 0.12 |
| LytTR-4CBV | (85, 99) | 0.05 | 0.05 |
| GerE-4E7P | (15, 104) | 0.05 | 0.05 |
| GerE-4E7P | (87 , 105) | 0.04 | 0.03 |

Table S7: Top ten interface predictions for the GerE query sequence using MSA Transformer. We compare to the average contact predictions using the OmpR and LytTR sequences.

| Interface contact subfamily | Interface contact | Contact prediction | Contact prediction OmpR-LytTR average |
|---|---|---|---|
| OmpR-1NXS | (90, 96) | 0.92 | 0.91 |
| GerE-4E7P | (6, 100) | 0.43 | 0.40 |
| GerE-4E7P | (6, 79) | 0.42 | 0.41 |
| OmpR-1NXS | (87, 109) | 0.20 | 0.20 |
| GerE-4E7P | (12, 104) | 0.20 | 0.20 |
| OmpR-1NXS | (91, 109) | 0.14 | 0.15 |
| OmpR-1NXS | (90, 109) | 0.12 | 0.12 |
| LytTR-4CBV | (85, 99) | 0.06 | 0.05 |
| GerE-4E7P | (15, 104) | 0.05 | 0.05 |
| GerE-4E7P | (87 , 105) | 0.05 | 0.03 |

sequences, sampling sequences from a probability distribution where the probability of selecting a sequence is inversely proportional to its number of neighbors ($\geq 80\%$ identity) in the MSA. To our knowledge, the MSAs we sample from are identical to those used to benchmark MSA Transformer in the ProteinGym study [29].

We provide a table of the results comparing MSA Pairformer to the ESM models and MSA Transformer below. Results for other are derived from the ProteinGym leaderboard.

Table S8: Average Spearman correlation between model scores and ProteinGym DMS experimental measurements. Experiments are grouped by coarse selection type. Bolded values indicate the largest in each category.

| | Avg. | Activity | Organismal Fitness | Binding | Expression | Stability |
|---|---|---|---|---|---|---|
| MSA Pairformer | **0.47 ± 0.16** | **0.49 ± 0.15** | **0.46 ± 0.14** | **0.35 ± 0.19** | 0.44 ± 0.15 | 0.51 ± 0.16 |
| MSA Transformer (ensemble) | 0.45 ± 0.17 | 0.48 ± 0.15 | 0.43 ± 0.17 | 0.32 ± 0.18 | **0.45 ± 0.15** | 0.49 ± 0.16 |
| ESM2 15B | 0.43 ± 0.18 | 0.42 ± 0.20 | 0.40 ± 0.17 | 0.31 ± 0.15 | 0.41 ± 0.17 | 0.49 ± 0.17 |
| ESM2 3B | 0.43 ± 0.19 | 0.43 ± 0.20 | 0.40 ± 0.19 | 0.31 ± 0.19 | 0.40 ± 0.18 | 0.51 ± 0.17 |
| ESM2 650M | 0.44 ± 0.20 | 0.44 ± 0.20 | 0.39 ± 0.21 | 0.33 ± 0.21 | 0.42 ± 0.17 | 0.52 ± 0.18 |
| ESM2 150M | 0.40 ± 0.22 | 0.40 ± 0.19 | 0.32 ± 0.22 | 0.32 ± 0.22 | 0.40 ± 0.15 | 0.51 ± 0.20 |
| ESM C 600M | 0.43 ± 0.21 | 0.43 ± 0.20 | 0.38 ± 0.22 | 0.29 ± 0.21 | 0.42 ± 0.18 | **0.53 ± 0.17** |
| ESM C 300M | 0.43 ± 0.21 | 0.43 ± 0.20 | 0.38 ± 0.22 | 0.31 ± 0.21 | 0.41 ± 0.18 | 0.53 ± 0.18 |

### B Additional references

[11] UniProt Consortium. Uniprot: a worldwide hub of protein knowledge. *Nucleic acids research*, 47(D1):D506–D515, 2019.

[12] UniProt Consortium. Uniprot: a hub for protein information. *Nucleic acids research*, 43(D1): D204–D212, 2015.

[13] UniProt Consortium. The universal protein resource (uniprot). *Nucleic acids research*, 36 (suppl_1):D190–D195, 2007.

[14] Allister Crow, Nicholas P Greene, Elise Kaplan, and Vassilis Koronakis. Structure and mechanotransmission mechanism of the macb abc transporter superfamily. *Proceedings of the National Academy of Sciences*, 114(47):12572–12577, 2017.

[15] Gustaf Ahdritz, Nazim Bouatta, Christina Floristean, Sachin Kadyan, Qinghui Xia, William Gerecke, Timothy J. O'Donnell, Daniel Berenberg, Ian Fisk, Niccolò Zanichelli, Bo Zhang, Arkadiusz Nowaczynski, Bei Wang, Marta M. Stepniewska-Dziubinska, Shang Zhang, Adegoke Ojewole, Murat Efe Guney, Stella Biderman, Andrew M. Watkins, Stephen Ra, Pablo Ribalta Lorenzo, Lucas Nivon, Brian Weitzner, Yih-En Andrew Ban, Shiyang Chen, Minjia Zhang,

Conglong Li, Shuaiwen Leon Song, Yuxiong He, Peter K. Sorger, Emad Mostaque, Zhao Zhang, Richard Bonneau, and Mohammed AlQuraishi. Openfold: retraining alphafold2 yields new insights into its learning mechanisms and capacity for generalization. *Nature Methods*, 21(8): 1514–1524, 2024.

[21] Mostafa Dehghani, Josip Djolonga, Basil Mustafa, Piotr Padlewski, Jonathan Heek, Justin Gilmer, Andreas Steiner, Mathilde Caron, Robert Geirhos, Ibrahim Alabdulmohsin, Rodolphe Jenatton, Lucas Beyer, Michael Tschannen, Anurag Arnab, Xiao Wang, Carlos Riquelme, Matthias Minderer, Joan Puigcerver, Utku Evci, Manoj Kumar, Sjoerd van Steenkiste, Gamaleldin F. Elsayed, Aravindh Mahendran, Fisher Yu, Avital Oliver, Fantine Huot, Jasmijn Bastings, Mark Patrick Collier, Alexey Gritsenko, Vighnesh Birodkar, Cristina Vasconcelos, Yi Tay, Thomas Mensink, Alexander Kolesnikov, Filip Pavetić, Dustin Tran, Thomas Kipf, Mario Lučić, Xiaohua Zhai, Daniel Keysers, Jeremiah Harmsen, and Neil Houlsby. Scaling vision transformers to 22 billion parameters, 2023. URL https://arxiv.org/abs/2302.05442.
